# Partially differentiated enterocytes in ileal and distal-colonic human F508del-CF-enteroids secrete fluid in response to forskolin and linaclotide

**DOI:** 10.1101/2025.02.03.636268

**Authors:** Yan Rong, Zixin Zhang, Hugo R. de Jonge, Ruxian Lin, Huimin Yu, Rafiq Sarker, Dario Boffelli, Rachel K. Zwick, Ophir D. Klein, Ming Tse, Mark Donowitz, Varsha Singh

## Abstract

Constipation causes significant morbidity in Cystic Fibrosis (CF) patients. Using CF patient (F508del) derived *ex vivo* ileal and distal colonic/rectal enteroids as a model and the Forskolin Induced Swelling Assay (FIS), we compared CFTR mediated fluid secretion in human enterocytes across the crypt-villus axis. <u>CFTR expression and FIS decreased as enterocytes differentiated from crypt to become partially differentiated and then mature villus cells</u>. While there was no FIS response in undifferentiated (crypt enterocytes) F508del-CF enteroids, partially differentiated F508del-CF enteroids had a swelling response to forskolin (cAMP) and linaclotide (cGMP) which was ∼48%, and ∼67% of the response in healthy enteroids, respectively and was prevented by a CFTR inhibitor. Also, linaclotide and a general PDE inhibitor independently enhanced combined CFTR-modulator-induced FIS response from partially differentiated F508del-CF enteroids. These findings demonstrate that partially differentiated ileal and distal colonic F508del-CFTR enteroids can be stimulated to secrete fluid by cAMP and cGMP.

## INTRODUCTION

The treatment of patients with Cystic Fibrosis (CF) with correctors/potentiators that target CFTR has led to life prolongation and reduced the requirements for major interventions such as lung transplantation ^1–3^. Consequently, additional morbidities associated with CF are becoming more frequently recognized and attempts to develop solutions have become a high priority. Chronic constipation is a significantly complication of CF which affects ∼47% of CF patients, despite appropriate treatment of pancreatic insufficiency, as well as concurrent CFTR modulator therapy ^4,5^. In CF patients, impaired anion (chloride (Cl^-^) and bicarbonate (HCO_3_) secretory defects reduce fluid secretion across many epithelia. Reduced anion secretion-driven intestinal water secretion as well as tenacious mucus are thought to be the major cause of constipation in CF. However, the constipation is contributed to by multiple additional factors that include, altered small intestinal or colonic <u>enteric nervous system</u> (ENS)/motor function, incompletely corrected malabsorption contributing to bulky stools, colonic inflammation, and changes in the microbiome ^6–8^. In addition, the accumulation of thick luminal mucus acts as a physical barrier to the absorption of some nutrients. Current treatments of constipation in patients with CF primarily include laxatives, enemas, and surgical intervention with no standard of care yet established ^9,10^.

Significant advances have been made in determining whether patients with specific CFTR mutations will respond to corrector/potentiator therapy by using an *ex vivo* functional assay based on Forskolin-Induced-Swelling (FIS) in patient-derived intestinal organoids or enteroids ^11–13^. These enteroids can be maintained in a crypt-like, undifferentiated state when grown in media containing growth factors Wnt3A, R-spondin, and Noggin, or they can be differentiated into villus-like or colonic surface cells by removal of Wnt3A and R-spondin for 5-6 days ^14–16^. In addition, we recently recognized that growth factor removal for 3-4 days resulted in an enterocyte population midway between the crypt-like Cl^-^ secretory cells and the well-differentiated villus or colonic surface enterocytes that take part exclusively in Na^+^ absorptive function ^14^. This represents a significant population of enterocytes that express transport proteins involved in NaCl absorption as well as anion secretion, including sodium-hydrogen exchanger-3 (SLC9A3, NHE3), chloride-bicarbonate exchanger-3 (SLC26A3, DRA), and CFTR. This population is called partially differentiated enterocytes, (abbreviated PD). We show in this study that ileal and distal colonic/rectal PD enteroids from the most common CFTR mutant-F508del, exhibit positive, CFTR-dependent fluid secretion in response to forskolin (cAMP) and the cGMP agonist linaclotide, unlike a lack of response in the undifferentiated CF enteroids. We further showed that linaclotide and a general phosphodiesterase inhibitor, theophylline independently enhanced CFTR-modulator-induced fluid secretion from PD enterocytes. Based on these pre-clinical studies, we suggest that strategies should be developed <u>to include ways to stimulate fluid</u> <u>secretion from</u> PD enterocytes in the treatment of constipation in CF patients.

## RESULTS

### Forskolin induced swelling (FIS) response is present in partially differentiated enteroids

The FIS response has been used to evaluate CFTR function by quantitating the rates of fluid secretion as rates of enteroid swelling ^11^. To define the FIS response in normal distal colonic/rectal <u>(to be referred to as colonic/rectal)</u> enteroids from <u>normal healthy subjects (NHS)</u> we compared fluid secretion in these enteroids grown in Wnt3A, R-spondin, and Noggin (undifferentiated (UD)), and the same enteroids grown in the absence of Wnt3A and R-spondin for 3 days /partially differentiated (PD)) and 6 days/fully differentiated (FD). Enteroid <u>cross-sectional surface area</u> was measured using live-cell microscopy, which showed a rapid expansion of both the lumen and total enteroid surface area in response to forskolin treatment, whereas DMSO-treated enteroids were minimally changed over the 1h study (Figure 1, A B). As shown in Figure 1, C-E, a forskolin stimulated, time-dependent surface area increase occurred in both UD (55.0 ± 6.0%, p<0.05 vs <u>DMSO</u>) and PD (36.0 ± 4.2% p<0.05 vs <u>DMSO</u>) NHS colonic/rectal enteroids at 60min, while FD enteroids showed no significant increase in the surface area (7.0± 4.2%) as compared to DMSO control. The FIS response in UD, PD, and FD enteroids correlates with the CFTR protein expression at different days of differentiation^14^ (Supplemental Figure 1A, B). To demonstrate that the FIS response was CFTR dependent and to evaluate the contribution of DRA, NHE3, and <u>Calcium-activated Chloride Channels</u> (CaCC), NHS colonic/rectal enteroids were preincubated for 1h with the CFTRinh-<u>Benzopyrimido-pyrrolo-oxazine-dione)</u> (BPO-27) (10µM), DRAinh A250 (10µM), NHE3inh S3226 (20 μM) or a general CaCCinh A01-(15µM) ^17–20^. Preincubation with BPO-27 significantly reduced FIS in both UD (10.4 ± 0.5%; p<0.05 vs Fsk) and PD (11.5 ± 0.5%; p<0.05 vs Fsk) NHS enteroids. In contrast, preincubation with the NHE3inh, DRAinh, and CaCCinh did not significantly affect the FIS response in either the UD or PD enteroids (Figure 1, C-E). This indicates no significant contribution of NHE3, DRA, or CaCC to the FIS assays in the normal colon/rectum. Taken together these results indicate that the FIS response in the colonic/rectal enteroids is primarily mediated by CFTR activity and the magnitude of the swelling response correlates with CFTR expression, both of which decrease with the enterocyte differentiation.

**Figure 1.**
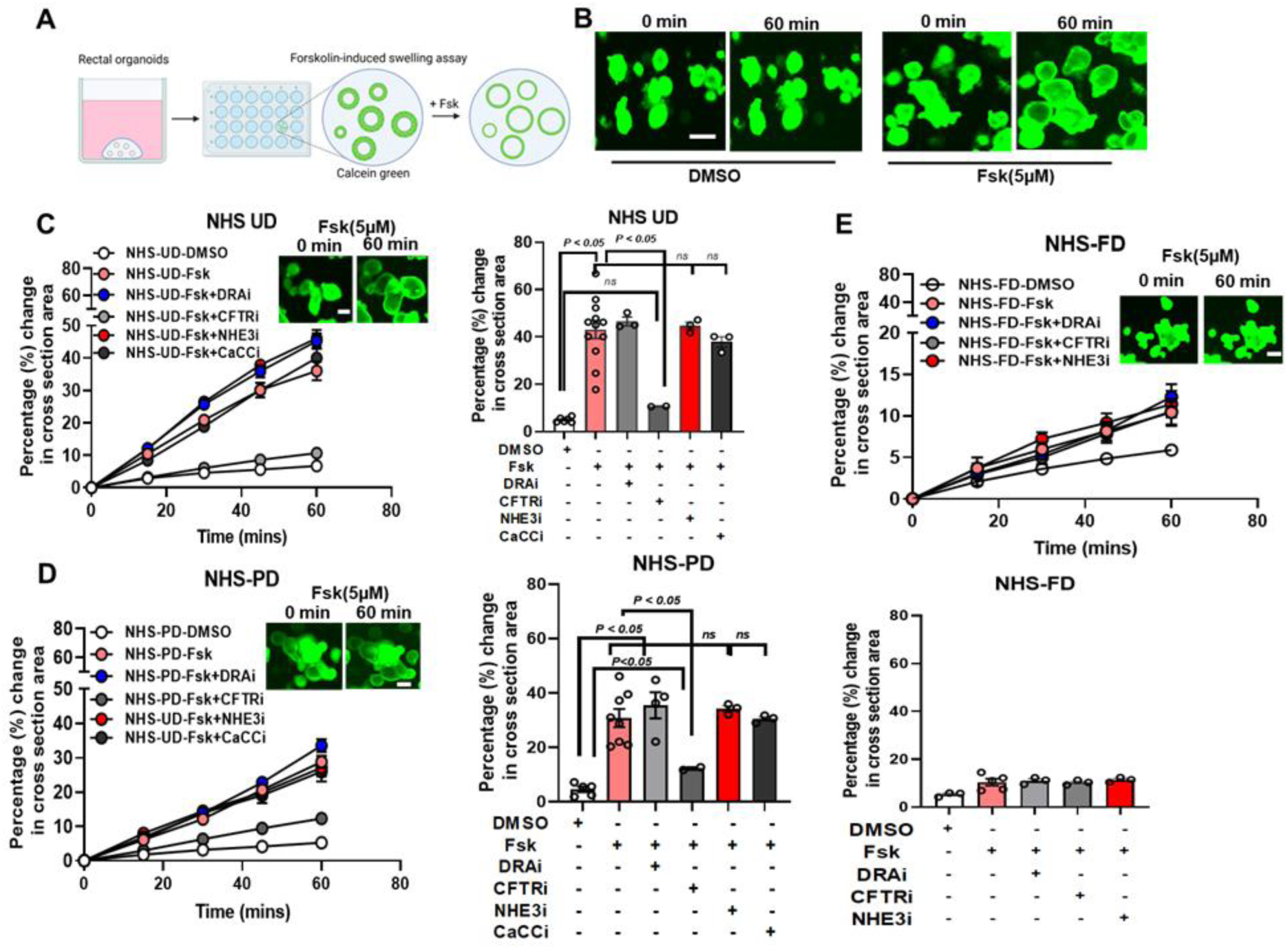
FIS response decreases but is present in partially differentiated enteroids. **(A)** Graphic illustration showing Forskolin-induced swelling (FIS) assay, and **(B)** representative fluorescence confocal images of a calcein green AM esters–labeled colonic/rectal enteroids from NHS treated with DMSO (control) or forskolin (Fsk, 5 μM). Images were taken at t = 0, and 60min after stimulation. Scale bar: 10µm. Quantification of FIS in NHS-UD **(C)**, PD **(D)** and FD **(E)**, alone or pretreated with CFTRinh BPO-27 (10µM), DRAinh A250 (10µM), NHE3inh S3226 (20 μM) or general CaCCinh-A01(15µM) for 1hr. Results are expressed as the percentage change in surface area relative to t = 0 (percentage change in cross-section area) measured at 15-min time intervals for 1 h (means ± SD), and the bar graph depicts fluid secretion plots (t = 60 min, means ± SEM, data points represent an average of 3-5 swelling assays in an individual donor, N=3 separate NHS). Analysis of differences was determined with a one-way ANOVA and Bonferroni post hoc test; ns=not significant.

### Partially differentiated F508del-CF enteroids have a positive CFTR-dependent FIS response

Previous studies in UD (undifferentiated) enteroids have established that CF enteroids do not respond to FIS unless the CFTR gene is corrected or treated with correctors/potentiators ^21–23^. Since CFTR was expressed in PD enteroids, we compared the FIS response in UD, PD, and FD (fully differentiated) NHS (normal healthy subject) and F508del-CF colonic/rectal enteroids. As reported, UD F508del-CF enteroid lacked a FIS response (6.5±1.1%). Unexpectedly, PD F508del-CF enteroids achieved a positive FIS response (12.7±1.3%), while FD enteroids had a small non-significant response (8.7±1.4%) (Figure 2, A). The magnitude of the FIS response in PD F508del-CF enteroids was approximately half (∼48±1.6%) of the response in PD NHS enteroids (Figure 2, B). This response was similar in colonic/rectal enteroids from three F508del-CF patients tested independently (Supplemental Figure 2A, B, and C). A similar positive FIS response was observed in PD ileal enteroids from a F508del-CF patient (UD:4.2±0.6%; PD: 12.6±0.4%; FD: 7.2±0.6%) (Supplemental Figure 3).

**Figure 2.**
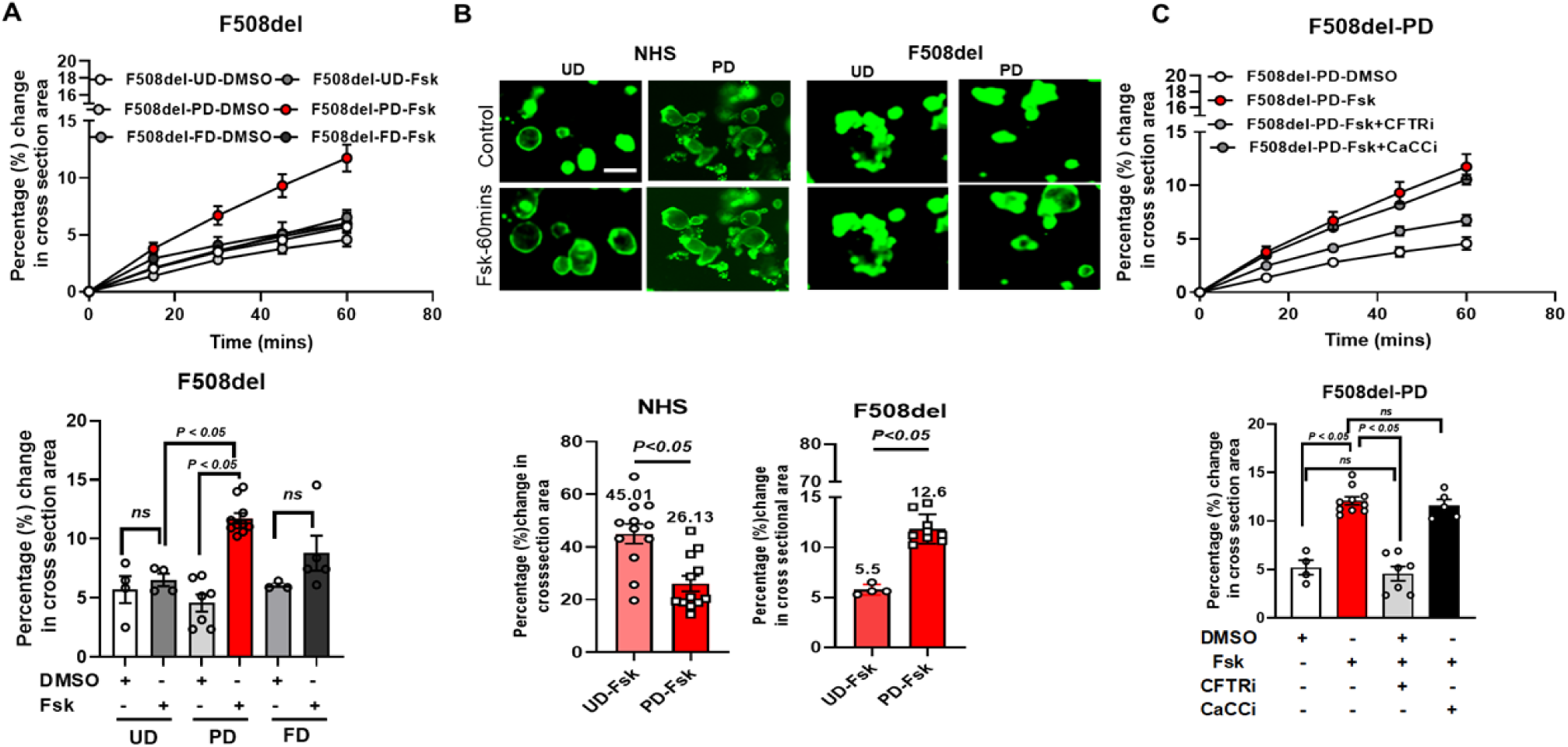
Partially differentiated F508del-CF colonic/rectal enteroids have a CFTR-dependent FIS response. **(A)** Quantification of FIS showing differences between UD, PD, and FD colonic/rectal enteroids from F508del-CF CF patients. Results are expressed as the percentage change in surface area relative to t = 0 (percentage change in cross-section area) measured at 15-min time intervals for 1 h (means ± SD), and the bar graph depicts fluid secretion plots (t = 60 min, means ± SEM). Data points represent an average of 5-6 independent assays each containing 50-100 enteroids from 1 CF patient (above). Combined results from all the 3 patients are shown (below). **(B)** A representative fluorescence confocal images (above) of a calcein green-AM esters–labeled enteroids from NHS and F508del-CF in UD and PD state treated with DMSO (control) or forskolin (Fsk, 5 μM) at t = 0, and 60min after stimulation. Below: bar graph of fluid secretion plots (t = 60 min, means ± SEM, data points represent an average of 10-15 independent assays from three NHS and F508del-CF CF patients). Analysis of differences was determined using a one-way Student’s t-test. Scale bar: 10µm. **(C)** Quantification of FIS in PD F508del-CF pretreated with CFTRinh BPO-27 (10µM), or CaCCinh-A01-(15µM) for 1hr or alone. Results are analyzed as above and expressed as means ± SEM. An average of 3-12 swelling assays each containing 50-100 enteroids from colonic/rectal enteroids from 3 separate F508del-CF patients is shown. Analysis of differences in (A) and (C) was determined with a one-way ANOVA and Bonferroni post hoc test, ns=not significant.

Since UD F508del-CF colonic/rectal enteroids were phenotypically different from NHS, enteroids we quantified the initial steady-state lumen area (SLA) of PD enteroids from both groups. This was done by comparing luminal area to total enteroid area in each group (Supplemental Figure 4, A and B) ^11^. For the colonic/rectal enteroids, there were no significant differences in the initial SLA between PD NHS and F508del-CF enteroids.

To identify the transport proteins contributing to the FIS response in PD F508del-CF colonic/rectal enteroids, the assay was performed in the presence of CFTRinh-BPO-27 (10µM) and CaCCinh-A01, (15µM). Like the response in PD NHS enteroids, the FIS response from PD F508del-CF enteroids was significantly inhibited by pretreatment with BPO-27, while A01 did not affect the FIS response (Figure 2, C). Thus, the positive FIS response in both NHS and F508del-CF PD enteroids is CFTR dependent and F508del-CFTR is secreting fluid in response to forskolin in this population. Due to these qualitatively similar FIS responses in ileal and colonic/rectal F508del-CF enteriods, results from colonic/rectal enteroids were reported for the rest of these studies.

To allow comparisons between the fluid secretory response between NHS and F508del-CF enteroids, differences in differentiation and cell types between F508del-CF and NHS enteroids, in UD, PD, and FD colonic/rectal enteroids were investigated. Compared to NHS, UD F508del-CF enteroids had significantly higher mRNA expression of Leucine-rich repeat-containing G-protein coupled receptor-5 (*LGR5*) and *Ki67,* used as a marker for proliferation, both of which gradually decreased with differentiation in PD and FD enteroids (Supplemental Figure 4C). The expression of the goblet cell marker *MUC2* mRNA increased with differentiation in PD and FD NHS enteroids, as well as in F508del-CF enteroids. In contrast, the expression of the enteroendocrine (EE) marker chromogranin A (*CHGA)* was significantly higher in PD and FD F508del-CF enteroids than in NHS enteroids (Supplemental Figure 4, C and D). Thus, F508del-CF enteroids had higher proliferative activity than NHS in the UD condition, and a higher number of CHGA-positive EE cells in FD enteroids but underwent differentiation that is similar to that in NHS enteroids.

The size of the PD enterocyte population has not been established. To estimate the size of this population, we further analyzed a published single-cell RNAseq dataset of normal human small intestine ^24^ using pseudotime description of CFTR, DRA, and NHE3 expressing cells. Pseudotime analysis showed that CFTR expression declined linearly going from stem cells to mature enterocytes; in contrast, expression of DRA and NHE3 was nearly absent in stem cells, <u>transit amplifying</u> (TA) cells, and enterocytes progenitors, increased in enterocytes and was maximal in mature enterocytes (Supplemental Figure 5). CFTR was significantly expressed in the area in which expression of DRA and NHE3 was approaching their maximum levels (pseudotime between 150 and 200). In this interval, the co-expression between these transporters were as follows: CFTR/DRA 88%; CFTR/NHE3 27%; DRA/NHE3 26%; CFTR/DRA/NHE3 25% (Supplemental Table 1 and Figure 5). Based on the pseudotime analysis, we propose that the cells at the transition zone from enterocytes to mature enterocytes likely represent PD enterocytes. This population represents 16% of villus enterocytes (calculated from cells in Supplementary Table 1, upper).

### Expression of DRA or inhibition of NHE activity do not contribute to fluid secretion in colonic/rectal enteroids

Previous studies in oocytes showed that SLC26 family members, SLC26A6 and SLC26A3, contribute to cAMP stimulation of CFTR activity ^25^. Since SLC26A3 is expressed in large amounts throughout the human colon and its expression significantly increases with differentiation^14,26^, we investigated whether DRA contributes to the FIS response in partially differentiated (PD) F508del-CF colonic/rectal enteroids. This was evaluated in two ways: 1) by overexpressing DRA in undifferentiated (UD) enteroids; and 2) by knocking down DRA and determining FIS in PD enteroids. To overexpress DRA, a replication-deficient adenovirus containing the coding sequence of human DRA (Flag-DRA), or containing no insert (control) were used to infect NHS and F508del-CF enteroids, followed by performing a FIS assay in the UD state. As shown in Figure 3A, Fsk (5µM) caused a similar FIS response in UD control and Flag-DRA transduced NHS-colonic/rectal enteroids (control: 59.3±2.0%; Flag-DRA: 62.6±0.7%). UD F508del-CF enteroids had no secretory response to Fsk even after being transduced with Flag-DRA. Western blot analysis of enteroids collected post-assay confirmed the expression of Flag-DRA in infected enteroids, while UD enteroids do not express DRA (Figure 3, A right).

**Figure 3.**
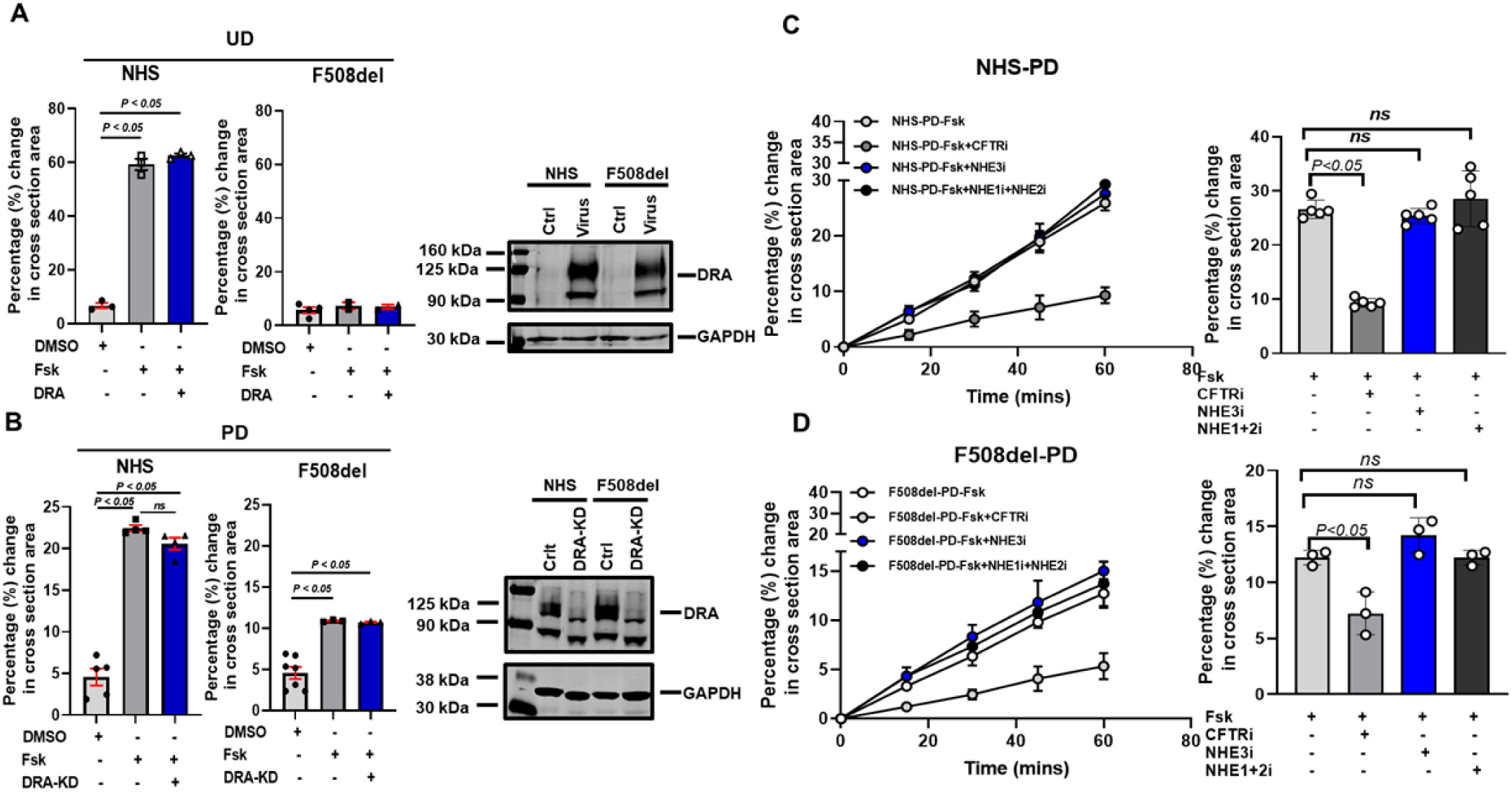
Expression of DRA or inhibition of NHEs does not affect fluid secretion in NHS and F508del-CF colonic/rectal enteroids. Bar graph showing quantitation of FIS response in colonic/rectal enteroids from **(A)** UD NHS and F508del-CF overexpressing control or DRA-adenovirus with representative western blot (right) of enteroids collected post-assay showing overexpression of Flag-DRA, and **(B)** PD NHS and F508del-CF enteroids overexpressing control or DRA-shRNA, with representative western blot (right) showing KD of DRA in DRA-shRNA (DRA-KD) expressing enteroids. Data are means ± SEM. Quantification of FIS in PD colonic/rectal enteroids from **(C)** NHS and **(D)** F508del-CF pretreated with BPO-27 (10µM), dimethyl amiloride (2µM), or S3226 (20μM). Results are expressed as the percentage change in the cross-sectional (surface) area relative to t = 0 measured at 15-min time intervals for 1 h (means ± SD), and the bar graph depicts fluid secretion plots (t = 60 min, means ± SEM). Data points represent an average of 3-5 independent assays each containing 50-100 enteroids from one NHS or F508del CF patient. Combined results from N=3 independent NHS and F508del-CF patients are shown. Analysis of differences in (A), (B), (C), and (D) was determined with a one-way ANOVA and Bonferroni’s post hoc test, ns=not significant.

Using lenti-shRNA against DRA or a scrambled sequence as control, a stable DRA knockdown was made in NHS and F508del-CF colonic/rectal enteroids. Western blot analysis of shRNA-infected cells showed reduced DRA protein expression by >40% in both NHS and F508del-CF enteroids compared to the control (Figure 3, B right). Furthermore, this reduction continued when these enteroids were maintained in a culture medium containing puromycin. These cells were used to measure the FIS response after partial differentiation. As shown in Figure 3B, PD colonic/rectal enteroids from both NHS and F508del-CF with reduced DRA expression had a significant FIS response (NHS: 22.2±1.5%; F508del-CF: 12.0±1.5%). <u>This response was not</u> <u>significantly different than the control-shRNA expressing enteroids, which were 20.2 ± 1.02%</u> <u>for NHS and 11.7 ± 1.5% for F508del-CF (</u>Figure 3B).

Partially differentiated colonic/rectal enteroids express a significant amount of NHE3 and these cells are also a major site for NHE2 expression ^27^. Inhibition of NHE3 results in enhanced fluid accumulation into the intestinal lumen due to the loss of the Na^+^ absorptive function of enterocytes ^24,28,29^. We hypothesized that the FIS response from PD F508del-CF enteroids could be due to cAMP (Fsk) induced inhibition of brush border NHE3 activity. Therefore, we measured the FIS in PD enteroids pretreated (1h) with the specific NHE3 inhibitor, 20µM S3226 (3-[2-(3-guanidino-2-methyl-3-oxo-propenyl)-5-methyl-phenyl]-*N*-isopropylidene-2-methyl-acrylamide dihydro-chloride), or with CFTRinh BPO-27 (10µM) ^19,30^. Pretreatment with 2 µM dimethyl amiloride (DMA) was used to inhibit NHE2 and NHE1 activities, to ensure the specificity of the contribution of NHE3. As shown in Figures 3, C, and D, the FIS response in NHS and F508del-CF colonic/rectal enteroids were not affected by exposure to DMA or S3226, compared to the FIS in their absence, while the CFTR inhibitor reduced the FIS response as in Figures 1 and 2. Overall, these results indicate that the FIS response in colonic/rectal enteroids does not depend upon the expression of DRA, NHE3, NHE2, or NHE1 activities.

### Linaclotide stimulates fluid secretion from partially differentiated (PD) colonic/rectal enteroids

Since PD F508del-CF colonic/rectal enteroids achieved a partial fluid secretion in response to cAMP (FSK) (Figure 2, A and B), we investigated the effect of another second messenger, cGMP on fluid secretion from PD enteroids. For this, we used the <u>guanylate cyclase-C</u> (GC-C) agonist linaclotide. Linaclotide stimulates the production and accumulation of intracellular cGMP, which acts as a second messenger to enhance fluid secretion in the intestine ^31,32^. The effect of linaclotide on CFTR regulation, particularly in human colonic/rectal samples, and the pathways associated with these effects have not been reported. The linaclotide (10µM) effects on the swelling response in UD, PD, and FD NHS, and F508del-CF colonic/rectal enteroids were determined over 2hrs. As shown in Figure 4, A and B, linaclotide did not cause a swelling response in UD NHS (4.5±2.1%) or F508del-CF (5.7±1.4%) enteroids. In PD and FD NHS enteroids, linaclotide alone (in the absence of forskolin) caused a time-dependent fluid secretion (PD: 18.3±1.3%; FD: 13±1.1%). The linaclotide-induced swelling from PD NHS enteroids was significantly higher than that from FD enteroids (p<0.05 vs PD). Similarly, PD F508del-CF enteroids, showed a significant swelling response to linaclotide (12.8±0.5%). Linaclotide did not cause a statistically significant swelling response in FD F508del-CF enteroids, compared to the untreated conditions (9.4±0.4% vs 13.9±2.2%; p=ns) (Figure 4, B). The swelling response to linaclotide in PD F508del-CF enteroids was ∼67% of the linaclotide response in PD NHS enteroids (Figure 4, A, and B).

**Figure 4.**
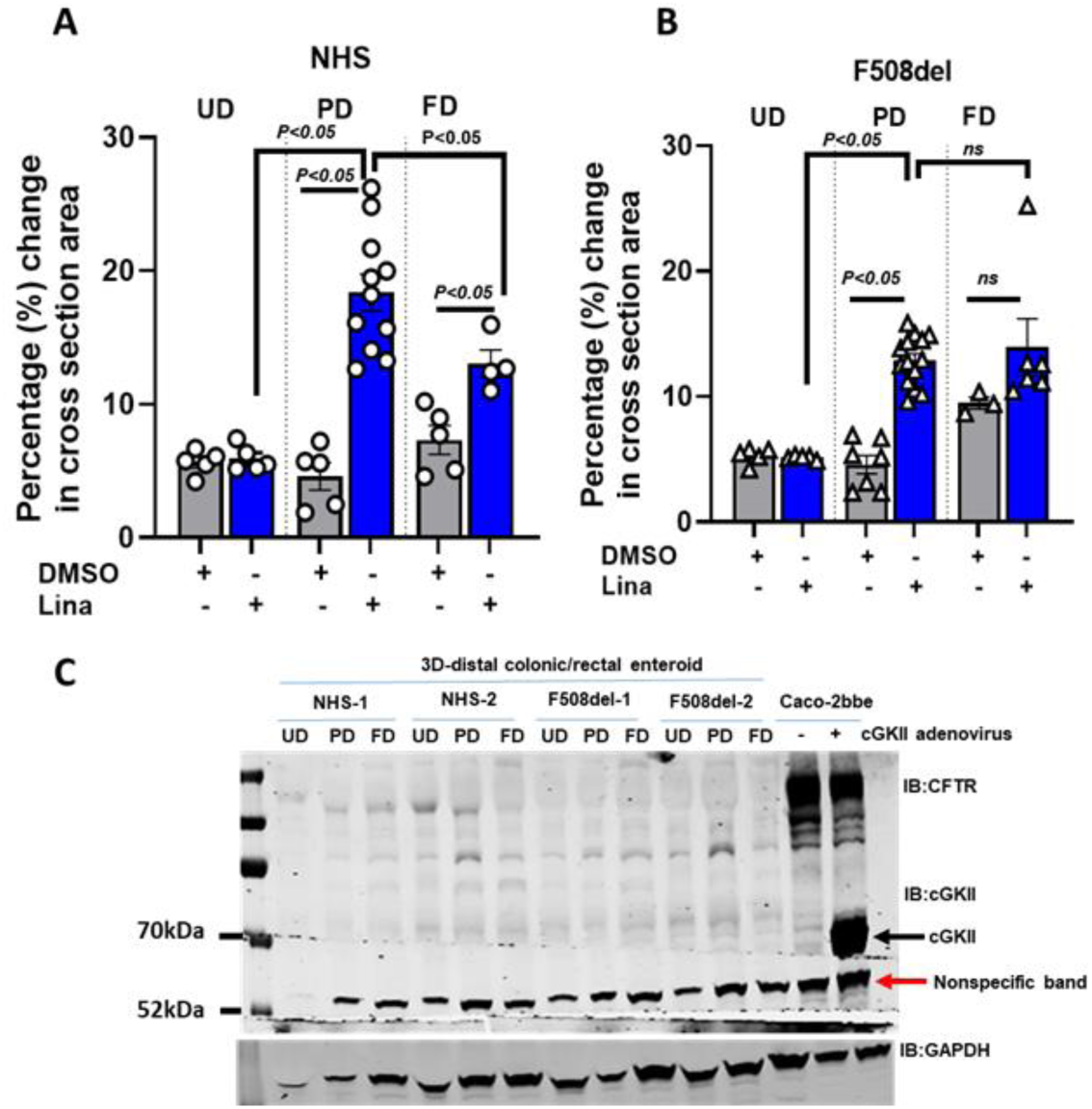
Linaclotide stimulates fluid secretion from partially differentiated colonic/rectal enteroids. Quantification of linaclotide (10µM) effect on swelling response in UD, PD, and FD colonic/rectal enteroids from **(A)** NHS and **(B)** F508del-CF patients measured over a 2h period. A time-dependent increase in organoid surface area normalized to that at t = 0 was calculated. Results are represented as a bar graph depicting fluid secretion plots (t = 120 min, means ± SEM). Data points represent an independent experiment consisting of 50-100 enteroids, N=3 separate NHS or F508del-CF. Analysis of differences was determined with a one-way ANOVA and Bonferroni post hoc test, ns=not significant. **(C)** Representative western blot from SDS-PAGE analysis of lysates prepared from UD, PD, and FD colonic/rectal enteroids from 2 separate NHS and F508del-CF patients and lysates from Caco2-bbe cells expressing control or cGKII-adenovirus as a positive control.

Linaclotide is known to employ the cGMP/cyclic-GMP-dependent kinase II (cGKII) pathway to elicit intestinal fluid secretion. To confirm that the essential signaling machinery is present, in addition to the presence of GCC (Supplemental Figure 4, C and D), we examined the cGKII expression in UD, PD, and FD colonic/rectal enteroids from NHS and F508del-CF patients. As a positive control, lysate from Caco-2/bbe cells transduced with adenovirus containing the coding sequence for rat cGKII was used ^33^. A 70kDa cGKII protein band was detected in lysates from Caco2-bbe-cGKII cells, but not from either NHS or F508del-CF enteroids, while a lower-sized non-specific band was present in lysates from NHS and F508del-CF enteroids as well as in Caco-2 cells not transduced with cGKII (Figure 4, C). Overall these results suggest that linaclotide stimulates fluid secretion from PD distal colonic/rectal enteroids that do not express significant amounts of cGKII.

### Linaclotide-stimulated fluid secretion in colonic/rectal enteroids is partially CFTR, and NHE3-dependent and involves additional transport processes

To determine the contribution of CFTR to linaclotide-induced fluid secretion, PD NHS and F508del-CF enteroids were preincubated with BPO-27 (10µM) for 1h before the linaclotide-induced swelling assay. Preincubation with BPO-27 partially reduced the linaclotide effect by ∼33% (18.8±1.4; 12.4±0.7) in NHS and ∼16% (13±0.4; 11±0.7) in F508del-CF PD enteroids (Figure 5, A, and B). This indicates that while CFTR contributes to linaclotide-mediated fluid secretion from PD NHS and F508del-CF distal colonic/rectal enterocytes, an additional transport protein(s) is involved. Previous reports suggest that linaclotide-induced gut fluidity is caused by inhibition of NHE3 activity ^34^. The way NHE3 inhibition could contribute to the linaclotide-induced swelling response would be by reducing the rate of fluid absorption from the enteroids in response to the cGMP increase. Thus to understand the role of NHE3 in the linaclotide response from PD enteroids, we pretreated the enteroids with NHE3_inh_ S3226 (20 μM) and then measured linaclotide-induced swelling. We observed a partial decrease in linaclotide response in the presence of S3226 in PD NHS as well as in F508del-CF distal colonic/rectal enteroids (NHS:14.4±1.1%; F508del-CF:10.6±0.7%) (Figure 5, C). We also investigated whether a combination of S3226 (20 μM) + BPO-27 (10µM) would further inhibit the linaclotide-induced swelling response. The pretreated combination of S3226 + BPO-27 further blocked the effect of linaclotide, but there was a significant residual linaclotide swelling response in both NHS (10.5±2.0%) and F508del-CF (8.8±1.03%) enteroids. This suggests the involvement of an additional transport protein(s) beyond NHE3 and CFTR in the linaclotide-induced swelling from PD enteroids in both NHS and F508del-CF distal colonic/rectal enteroids.

**Figure 5.**
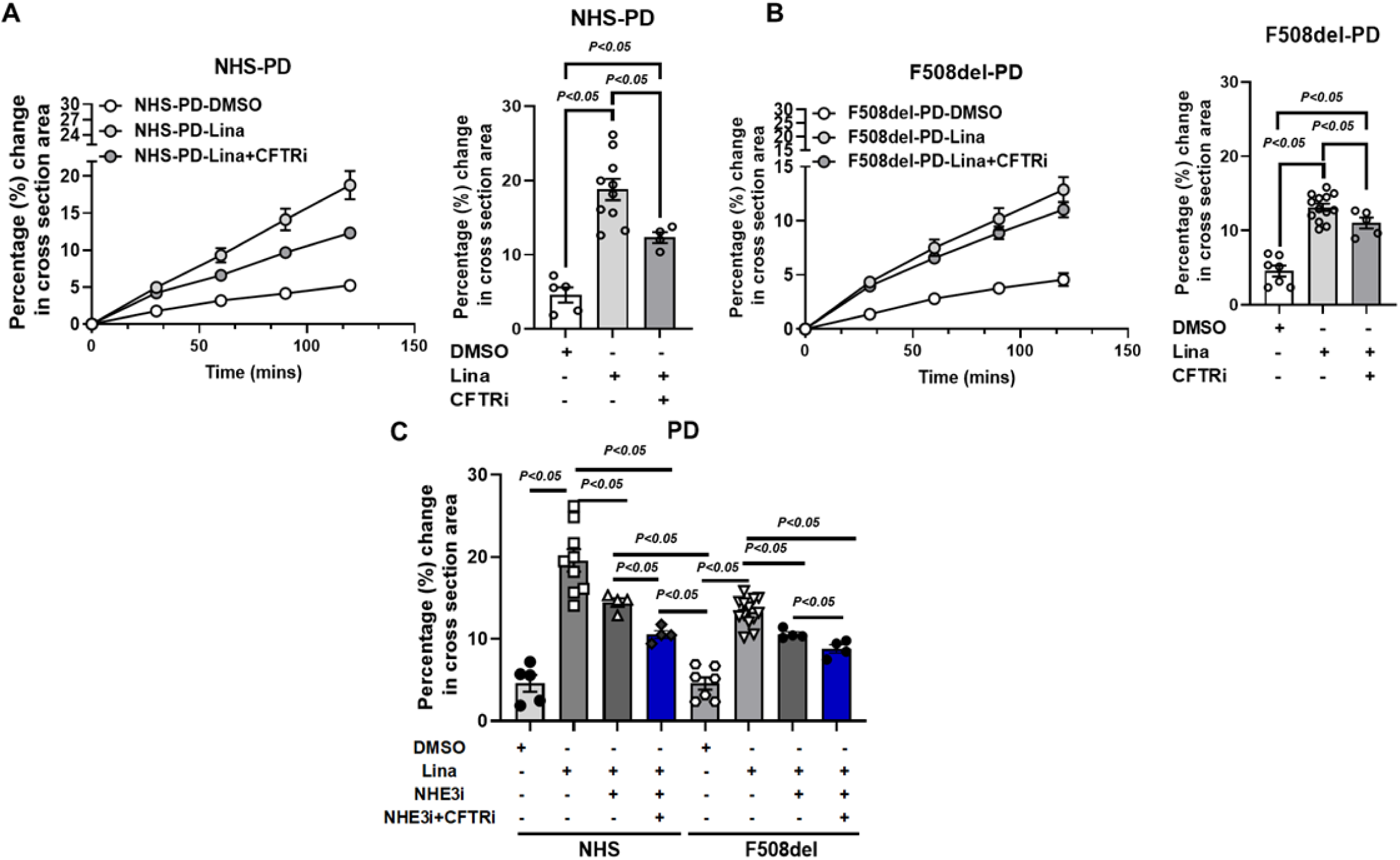
Linaclotide-stimulated fluid secretion in colonic/rectal enteroids was partially blocked by CFTRinh-BPO, and NHE3inh S3226. The role of CFTR in Linaclotide (10µM)-induced swelling was measured by pretreating PD enteroids from **(B)** NHS and **(D)** F508del-CF patients with BPO-27 (10µM). A time-dependent increase in enteroids surface area normalized to that at t = 0 was measured at 15-min time intervals for 2 h (means ± SD). Bar graph depicting fluid secretion plots (t = 120 min, means ± SEM). Data points represent an independent experiment consisting of 50-100 enteroids, N=3 separate NHS and F508del-CF. Data points represent an independent experiment consisting of 50-100 enteroids, N=3 separate NHS and F508del-CF. Analysis of differences in (A) and (B) were determined with a one-way ANOVA and Bonferroni post hoc test. (**C)** Linaclotide-induced swelling in PD NHS (left) and F508del-CF distal colonic/rectal enteroids (right) was performed alone or after pretreatment with S3226 (20 μM) or S3226 (20 μM) + BPO-27 (10µM) for 1hr. An increase in enteroid surface area in response to Linaclotide (10µM) normalized to that at t = 0 was analyzed as above and expressed a bar graph depicting fluid secretion plots (t = 120 min, means ± SEM). Data points represent an average of 2-3 independent assays in enteroids from NHS and F508del-CF patients, N=3 separate NHS and F508del-CF. *P* value was determined by 1-way ANOVA with Bonferroni’s adjustment for multiple comparisons.

### F508del-CF treatment with combined CF correctors (elexacaftor+tezacaftor) and a potentiator (ivacaftor) partially increases CFTR-mediated fluid secretion from UD and PD colonic/rectal enteroids

Since CFTR correctors and a potentiator increase the activity of F508del-CFTR, the response of these CF drugs in undifferentiated (UD) and partially differentiated (PD) F508del-CF colonic/rectal enteroids was determined. For these studies, the enteroids were exposed to CF correctors: elexacaftor+tezacaftor 3µM each for 24h and the potentiator: ivacaftor 5µM for 1h. This is the combination of corrector and potentiator approved by the FDA as triple therapy in Trikafta ^35,36^. We refer to this combination as <u>combined CFTR-modulators</u>. Undifferentiated and PD F508del-CF enteroids were pretreated with elexacaftor+tezacaftor 3µM each for 24h and with ivacaftor for an hour and FIS was measured and compared with DMSO-treated control enteroids; in addition the effects of DRA_inh_ A250 (10µM) and CFTR_inh_ BPO-27 (10µM) were also determined. As shown in Figure 6, A, UD F508del-CF enteroids had a positive FIS response only after treatment with <u>combined CFTR-modulators</u> (27.1±2.0%). This FIS response was less than the response in UD NHS enteroids performed in the absence of a <u>combined CFTR-modulators</u> (43±3.8%, Figure 1, C). The Fsk response in <u>combined CFTR-modulator</u> treated F508del-CF enteroids was reversed by pretreatment with the CFTRinh (8.1±1.0%) but not by the DRAinh (Figure 6, A). A similar assay was performed in PD F508del-CF enteroids. The PD F508del-CF enteroids without CF correcting drugs had a FIS response of 11.9±0.4%; when treated with the <u>combined CFTR-modulators</u>, the FIS response was enhanced to 18.1±1.5%, p<0.05. As in UD enteroids, significant FIS response was lost with pretreatment with the CFTR inhibitor but not with the DRAinh (Figure 6, B). Together these results suggest that the correctors/potentiator that make up Trikafta increase CFTR function from UD and PD F508del-CFTR enteroids. F508del-CFTR is partially activated by the cAMP in PD enteroids and the addition of <u>combined CFTR-modulators</u> can further enhance this activity.

**Figure 6.**
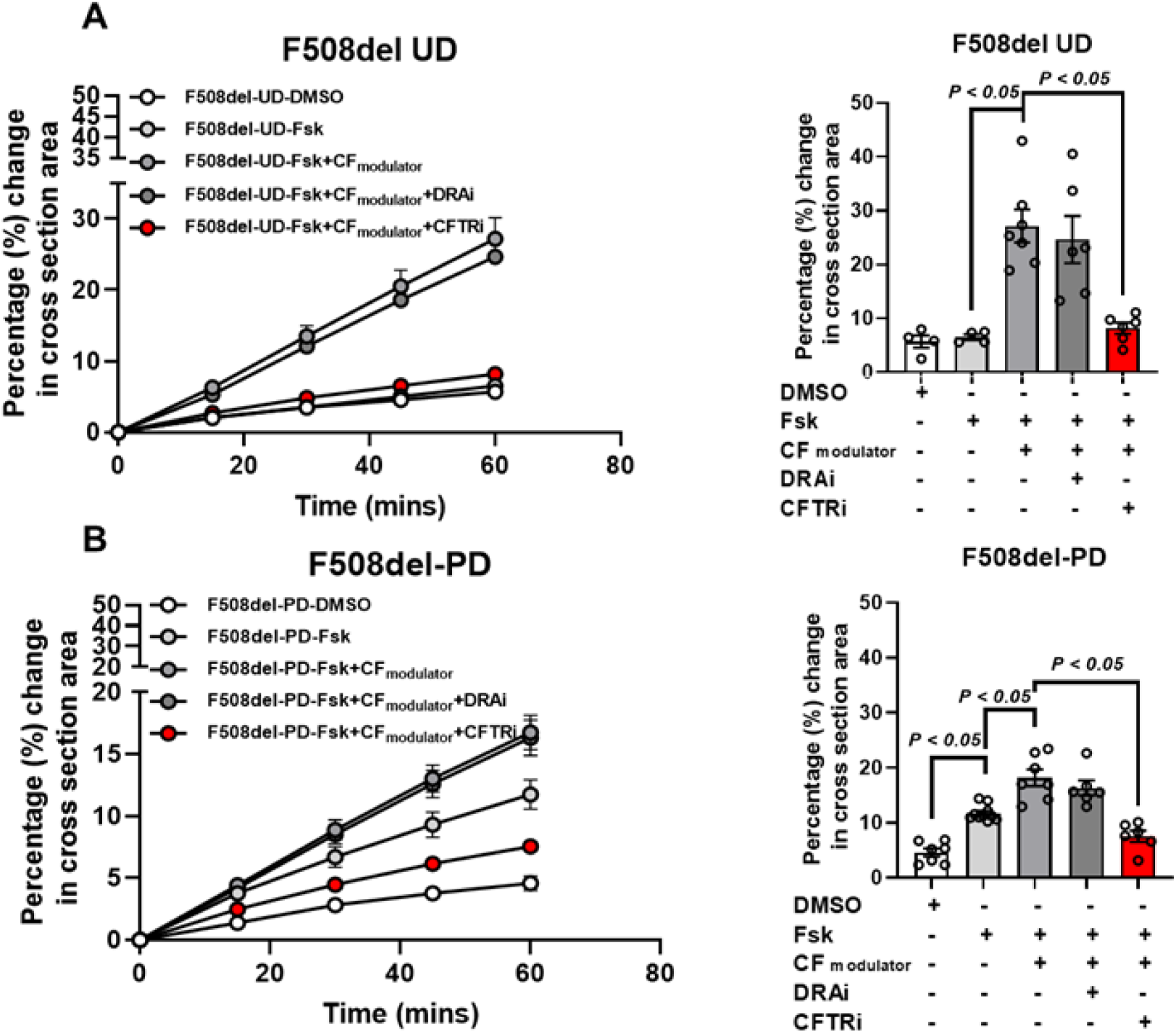
F508del-CF correction with combined CFTR-modulators enhances CFTR-mediated fluid secretion from UD and PD colonic/rectal enteroids. FIS in **(A)** UD and **(B)** PD F508del-CF colonic/rectal enteroids under basal or in response to combined CFTR-modulators (tezacaftor:VX-661+elexacaftor:VX-445 3µM each for 24h followed by ivacaftor:VX-770 (5µM) for 1h) pretreated with vehicle or BPO-27 (10µM) or A250 (10µM). Untreated or combined CFTR-modulators-treated colonic/rectal enteroids were stimulated with Fsk (5µM), CFTR_inh_ BPO-27, and DRA_inh_ A250. Results are expressed as the percentage change in surface area relative to t = 0 measured at 15-min time intervals for 1 h (means ± SD), and the bar graph depicts fluid secretion plots (t = 60 min, means ± SEM). An average of 3-12 swelling assays each containing 50-100 enteroids from distal colonic/rectal enteroids from N=3 separate F508del-CF patients is shown. Analysis of differences was determined with a one-way ANOVA and Bonferroni post hoc test.

### A non-selective phosphodiesterase inhibitor, theophylline, and linaclotide independently enhance the response of the combined CFTR-modulators on fluid secretion in F508del-CF colonic/rectal enteroids

The effect of the <u>combined CFTR-modulators</u> on the FIS response from undifferentiated (UD) and partially differentiated (PD) distal colonic/rectal F508del-CF enteroids was lower than the FIS response in untreated NHS enteroids. To maximize the effect of the <u>combined CFTR-modulators</u>, we investigated the effect of two pharmacological compounds: i) a non-selective phosphodiesterase (PDE) inhibitor, theophylline, and ii) linaclotide, on the <u>combined CFTR-modulators</u> effect on fluid secretion. Phosphodiesterases (PDEs) are a group of enzymes that catalyze the hydrolysis of phosphodiester bonds of the second messengers, cAMP, and cGMP, thus reducing the cyclic nucleotide levels ^37,38^. Inhibition of PDEs has been used to stimulate CFTR activity and to treat patients with asthma and chronic obstructive pulmonary disease ^39^. Colonic/rectal enteroids were pretreated with theophylline (100µM-1h) which was present throughout the assay period. As shown in Figure 7, A, the FIS response in UD NHS enteroids was significantly enhanced due to PDEinh (Fsk:45±3.7; Fsk+PDEinh: Fsk:65±2.1; p<0.05). In UD F508del-CF enteroids, theophylline exposure did not cause a FIS response, unless CF enteroids were exposed to the <u>combined CFTR-modulators</u> (24.5±2.0%), which allowed theophylline to significantly enhance the fluid secretion (37.6±1.1%; p<0.05) (Figure 7, B). Similarly, in PD F508del-CF enteroids the FIS response was not significantly affected by theophylline treatment (13.3±5.1%; 11.9±5.1%; NS); however, the <u>combined CFTR-modulators</u> increased FIS was significantly enhanced by theophylline treatment (16.3±1.4%; 22.4±2.1%;p<0.05) (Figure 7, B). Colonic/rectal enteroids from all three F508del-CF patients had a similar theophylline-related enhancement of the combined CFTR-modulators stimulated secretion (Supplemental Figure 6). Of note, F508del-CF colonic/rectal enteroids had significantly higher mRNA expression of PDE (PDE4) than NHS (Supplemental Figure 4, C).

**Figure 7.**
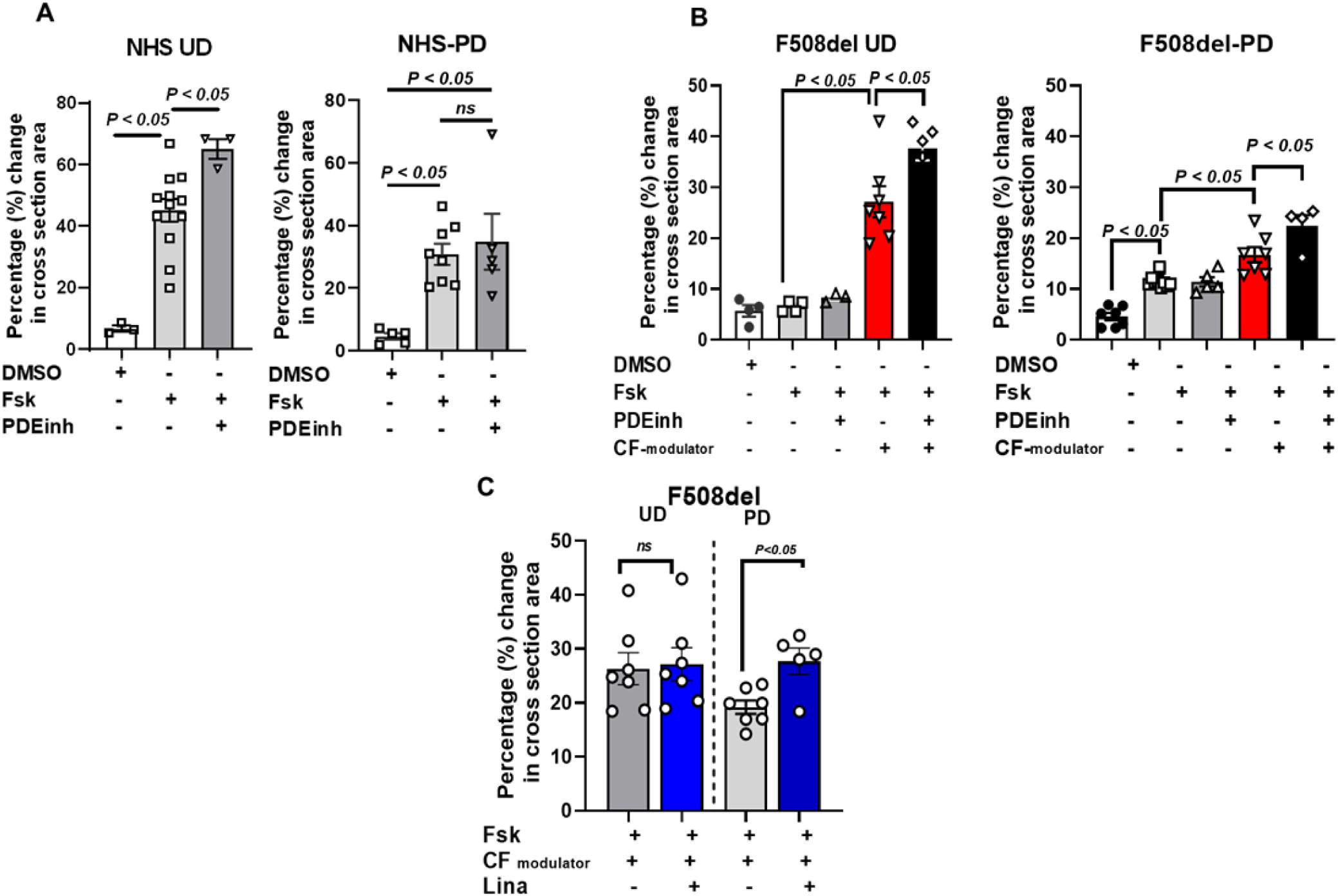
Theophylline and linaclotide independently enhance the combined CFTR-modulators response from F508del-CF colonic/rectal enteroid. **(A)** Effect of theophylline (100µM-1h) on FIS (5µM) response in UD, and PD enteroid from NHS. **(B)** Effect of Theophylline (100µM-1h) or combined CFTR-modulators (tezacaftor +elexacaftor (3µM) for 24h; ivacaftor (5µM) for 1h) and in combination on FIS (5µM) in UD, and PD F508del-CF enteroid. Changes in surface area relative to t = 0 were calculated and the bar graph depicts fluid secretion plots (t = 60 min, means ± SEM). Data points represent an independent experiment consisting of 50-100 enteroid, n=3 separate NHS. **(C)** Quantitation of Linaclotide effect on FIS response in combined CFTR-modulators treated UD and PD F508del-CF enteroids. Enteroids were pretreated with combined CFTR-modulators as above and were used to determine FIS in the presence of DMSO or Linaclotide (10µM). Results were analyzed as above and expressed as means ± SEM of fluid secretion plot at t=120 mins. Data points represent an average of 2-3 independent assays in enteroid from NHS and F508del-CF patients, n=3 separate NHS and F508del-CF. *P* value by 1-way ANOVA with Bonferroni’s adjustment for multiple comparisons.

For the linaclotide response, the fluid secretion was measured in the combined CFTR-modulators treated UD and PD colonic/rectal enteroids from F508del-CF patients in response to Fsk alone or with linaclotide (10µM) over a 2hr period. As shown in Figure 7, C, UD and PD F508del-CF colonic/rectal enteroids treated with the <u>combined CFTR-modulators</u> had significant fluid secretion in response to Fsk (UD:26.3±2.9%; PD:19.2±1.3). Linaclotide further enhanced this fluid secretion but only in PD F508del-CF enteroids (Fsk+combined CFTR-modulators:19.2±1.3%; Fsk+<u>combined CFTR-modulators</u>+linaclotide: PD:27.7±2.5%, p<0.05). Linaclotide enhanced the fluid response in PD enteroids in each of the three F508del-CF patients (Supplemental Figure 7). These results suggest that inhibition of PDE and the addition of linaclotide should be considered for testing as a strategy to enhance the fluid secretory effect of the <u>combined CFTR-modulators</u> in F508del CF patients.

## DISCUSSION

Constipation in CF patients is multifactorial. However, given that the primary defect in CF relates to reduced anion secretion, an accepted driver of stool water amount, most emphasis in approaching ways to deal with CF patient constipation has been to improve the reduced anion secretion. Formal clinical trials of treatment for constipation in CF patients are limited. In a study of 438 CF patients on some combinations of elexacaftor/ tezacaftor/ ivcaftor, for up to 6 months, there was no clinically significant changes in bowel characteristics, symptoms related to chronic constipation, or constipation-related quality of life assessments ^40^. Despite no evidence demonstrating efficacy, current approaches to treat CF constipation include the use any of a multitude of laxatives including lubiprostone, linaclotide as well as osmotic agents, and take advantage of the demonstration that CFTR correctors plus potentiators enhance CFTR-mediated anion secretion in most CF patients ^35,41^. Based on our findings, we suggest that an effective strategy for enhancing the treatment of chronic constipation in CF patients is to utilize a combination of agonists, which alone or together, enhance the increased fluid secretion caused by the correctors and potentiators, while also targeting the enterocyte populations studied in this research.

Using human enteroids as a preclinical model, we showed that CFTR is the major contributor to the fluid secretion in the distal colonic/rectal enterocytes and its expression and secretory activity decreases as the enterocytes move from crypts to become differentiated villus/surface cells. In addition, we previously showed that as the crypt cells differentiate they are called partially differentiated enterocytes, which express transport proteins, have and characteristics of both the crypt and mature villus ^14^. Here we demonstrate that in the small intestine, these cells express CFTR together with DRA and often with NHE3. This population of cells secrete fluid in response to forskolin (cAMP), although less in amount compared to crypt cells. The percentage of the PD enterocyte population in the human small intestine and colon has not yet been established. The previously reported scRNA-seq analysis of the human small intestine has suggested that there is a significant population of villus enterocytes expressing DRA and CFTR (Supplemental Table 1). This result is supported by the reanalysis of a recent human ileal single-cell-RNAseq report including privileged information provided by *Magness et al.,* that divided ileal enterocytes into early (∼UD), intermediate (∼PD), and late (∼FD) populations ^42^. This result showed that a significant number of the intermediate enterocyte populations express DRA, CFTR, and NHE3. Nonetheless, the quantitative contribution of the PD enterocytes awaits single-cell proteomic analysis. Importantly, NHE3, DRA, and CFTR when present in the same cells (as in PD enterocytes), interact dynamically, supporting that there appears to be unique regulation, particularly of DRA in this population ^14,43^.

We used the FIS assay developed by Dekkers *et al* to evaluate the quantitative contribution of the PD enterocytes in comparison with the UD (crypt) and FD (villus) enterocyte populations ^44^. Using human colonic/rectal enteroids we showed that F508del-CF UD enteroids had a positive FIS response only following treatment with the <u>combined CFTR-modulators</u> and with the non-selective phosphodiesterase inhibitor theophylline. In contrast, PD F508del-CF enteroids had a positive FIS response to increased cAMP alone, which was further enhanced by theophylline, and by the <u>combined CFTR-modulators</u>, and there was increased secretion when the <u>combined</u> <u>CFTR-modulators</u> were studied together with theophylline. Additionally, PD F508del-CF enteroids secreted fluid in response to linaclotide and there was a larger effect of the <u>combined</u> <u>CFTR-modulators</u> in the presence of linaclotide. Thus, this PD enterocyte population appears to represent a potentially important, previously unrecognized drug target for increasing intestinal (ileum and distal colon/rectal) fluid secretion in CF patients. The additivity of the effect of theophylline to the FIS assay and the correctors+potentiator response suggests that the assay is not saturating under the conditions of the study and that multiple agonists used together may be useful in the treatment of CF constipation.

Linaclotide is emerging as a promising candidate for constipation in CF patients. However, no clinical studies of its efficacy have been reported on this patient population. Moreover, linaclotide use in treating otherwise healthy patients with chronic constipation is not effective in as many as 50% of patients ^45,46^. As previously established, linaclotide causes its effects by binding to guanylate cyclase C (GCC) and by increasing cGMP. Of note, intestinal GCC expression varies widely based on species differences; there is significant mRNA expression in mouse colonic crypts, but as demonstrated in this study, not in human colonic crypts. Moreover, it was reported in the initial molecular identification of cGKII, that rat transverse colon, distal colon, and rectum have very little cGKII, an observation shown here to be true in human colonic/rectal enteroids ^47^. In this context, it is surprising that linaclotide caused fluid secretion in PD enteroids in both NHS and F508del-CF colonic/rectal enteroids which lacked cGKII even though they did express GCC (Supplemental Figure 3C). While not tested in this study, high concentrations of cGMP have been reported to activate CFTR via PKA in Xenopus oocytes that lack cGKII ^48,49^.

The complex signaling for linaclotide is further indicated by reported studies of effects in (a) rat proximal colon, and (b) mouse jejunum, both of which express cGKII. In rat proximal colon, the linaclotide stimulated Cl^-^ secretion was partially inhibited by the cGKII inhibitor H8 but no studies probed what mediated the residual component, although, suggestions were made regarding the potential involvement of cGMP activation of PKA or inhibition of a phosphodiesterase inhibitor ^50,51^. To our knowledge, ours is the first report showing that linaclotide-induced fluid secretion occurs in the human distal colon/rectum and that this occurs specifically in the PD enterocyte populations. Of all the transport proteins investigated in the swelling assays of this study, CFTR contributed solely or partially to the swelling response, and NHE3 was the only other protein identified that contributed to the linaclotide-induced secretion. Importantly, a part of the linaclotide swelling response appeared to be independent of NHE3 plus CFTR, indicating the involvement of additional transport protein(s).

Our studies using normal human and F508del-CF enteroids add to the preclinical studies of ways to increase intestinal fluid secretion, which has been primarily carried out in mouse and rat intestines. The recognition that PD enterocytes represent a previously unrecognized enterocyte population with a unique distribution of transporters involved in Na^+^ and Cl^-^ transport and with unique acute regulation offers the opportunity to extend preclinical studies to understand ways to alter intestinal transport in CF patients to increase intestinal fluid secretion. While the percentage of the PD enterocyte population remains under evaluation, the magnitude of the FIS response in PD F508del-CF colonic/rectal enteroids with <u>combined CFTR-modulators</u> was ∼60% of that in UD F508del enteroids, even in the face of reduced CFTR expression. What accounts for the CFTR-dependent fluid secretion from PD F508del-CF enteroids is not known, but it does not appear to involve increased brush border expression of the CFTR since the C band was not increased in the PD compared to the UD F508del-CF enteroids. Further studies using single-cell RNAseq and proteomics from UD and PD F508del-CF distal colonic/rectal enterocytes are needed to identify which signaling molecules are unique to the PD population. Importantly, given that constipation in CF patients is multifactorial, clinical studies in patients treated with <u>combined CFTR-modulators</u> such as Trikafta combined with additional agents (cAMP, and cGMP agonists, and PDEinh) to treat constipation will be required to define the most helpful regimes, which may vary between individuals as well as based on the nature of the CF mutation.

### Limitations of the study

The size of the PD population in the human small intestine and colon has not yet been adequately defined. Single-cell RNAseq analysis of the human colon has not been reported as yet in parallel to the studies in the human small intestine shown in supplementary Fig 5 and reported in detail^52^. However, the presence of a partially differentiated colonic enterocyte population is supported by the similar maturation pattern of enterocytes from the small intestine and colon, with a similar although slightly longer half-life of the colonic enterocytes to approximately 5-7 days during which the crypt cells mature into surface enterocytes^53^. Importantly, while approximately two-thirds of normal human proximal colon in the Turner Laboratory Human Transport Atlas shows both CFTR and DRA in colonocytes in the upper crypt and/or surface cells, the cells expressing DRA with and without CFTR were intermingled. Thus, as in the past when we used undifferentiated vs fully differentiated human enteroids grown by altering the presence of growth factors (Wnt, R-spondin) to model and allow study separately of the intestinal crypt and villus^54^, we support that we are similarly able to use the altered presence of growth factors to study separately undifferentiated, fully differentiated and partially differentiated colonocytes.

For this study, the F508del-CFTR mutant was chosen because this accounts for about 70% of CFTR mutations worldwide and is present in approximately 85% of CF patients ^55,56^. Future studies are needed to understand if the PD enterocyte population has a similar active role in mutations in the other classes of defective CFTRs, which would indicate that the proposed pharmacological approach might be more generally useful for treating the constipation of CF patients.

Although enteroids demonstrate various cell types from their native tissue, they still exclusively contain epithelial cells. Thus, co-culture studies of intestinal enteroids with cell types such as ENS, endothelial, and immune cells are needed to examine the contribution of these cells to fluid secretion by interacting with epithelial cells. As of now, the co-culture systems remain relatively simple, incorporating only one or a few cell types, but failing to simulate the complexity of the intact intestine.

## Resource Availability

### Lead Contact

Further information and requests for resources and reagents should be directed to and will be fulfilled by the lead contact, Varsha Singh.

### Materials Availability

Enteroids generated in this study are available upon request as per Johns Hopkins University policy.

### Data and code availability

Previously published single-cell sequencing data^52^ analyzed here are available under accession code GSE201859.

The original dataset of immunofluorescence images and western blots are available upon reasonable request through the lead contact.

## Acknowledgments

The authors wish to acknowledge the P30-DK-89502, Hopkins Conte Digestive Disease Basic and Translational Research Core Center Integrated Physiology Core for enteroid and enteroid supplies, and the Imaging Core for IF studies. **Funding:** This study is partly supported by Cystic Fibrosis Foundation grant CF-SINGH20G0, National Institutes of Health grants R01-DK-116352 and P30-DK-89502 (the Hopkins Basic and Translational Research Digestive Diseases Research Core Center).

## Author Contributions

Conceptualization, V.S; methodology, Y.R., Z.Z., R.R., R.S., M.T.; resources, H.R., H.Y.; investigation, D.V., R.Z., O.K.; writing-original draft, V.S.; writing-review & editing, V.S, M.D.; funding acquisition, V.S, M.D.; supervision, V.S, M.D.

## KEY RESOURCES TABLE

| REAGENT OR RESOURCE | SOURCE | IDENTIFIER |
| --- | --- | --- |
| Antibodies |  |  |
| DRA | Santa Cruz | SC-376187 |
| cGKII | Santa Cruz | SC-393126 |
| GAPDH | Sigma-Aldrich | G8795 |
| CFTR | 660, 217, 596 | CF consortium |
| CHGA | Immunostar | 20085 |
| GUCY2C | LSBio | LS-C498735, |
| MUC2 | Santa Cruz | sc7314 |
| E-cadherin | BD Biosciences | 610182 |
| Goat anti-Rabbit IgG (H+L) Cross-Adsorbed Secondary Antibody, Alexa Fluor 488 | Life Technologies | A11008 |
| Goat anti-Mouse IgG (H+L) Cross-Adsorbed Secondary Antibody, Alexa Fluor 546 | Life Technologies | A11003 |
| Bacterial and virus strains |  |  |
| pLKO-1-DRA | addgene | 8453 |
| Biological samples |  |  |
| Chemicals, peptides, and recombinant proteins |  |  |
| Y-27632 | Selleck | S1049 |
| EGF Recombinant Mouse Protein | Invitrogen | PMG8043 |
| CHIR99021 | Selleck | S1263 |
| N-2 Supplement | Invitrogen | 17502-048 |
| B-27 Supplement | Invitrogen | 17504-044 |
| HEPES | Invitrogen | 15630-080 |
| GlutaMAX | Invitrogen | 35050-061 |
| CHIR99021 | Selleck | S1263 |
| Triton-X-100 | Sigma-Aldrich | 93443 |
| Matrigel | Corning | 356231 |
| Paraformaldehyde | Electron<br>Microscopy<br>Sciences | 15710 |
| Hoechst 33342 | Invitrogen | H3570 |
| Organoid Harvesting Solution | R&D System | 3700-100-01 |
| Acetylcysteine | Sigma-Aldrich | A7250 |
| Human epidermal growth factor | ThermoFisher | A42555 |
| Gastrin | AnaSpec | AS-64149 |
| A83-01 | Tocris | TB2939-RMU |
| SB202190 | MilliporeSigma | 152121-30-7 |
| Primocin | InvivoGen | ant-pm-05 |
| Critical commercial assays |  |  |
| Deposited data |  |  |
| Experimental models: Cell lines |  |  |
| Caco-2bbe | ATCC |  |
| Experimental models: Organisms/strains |  |  |
| Oligonucleotides |  |  |
| *LGR5* | ORIGENE | HP207145 |
| *Ki67* | ORIGENE | HP206104 |
| *MUC2* | ORIGENE | HP206138 |
| *CHGA* | ORIGENE | HP205193 |
| *DRA* | ORIGENE | HP200096 |
| *SI* | ORIGENE | HP203589 |
| *PDE4* | ORIGENE | HP233178 |
| *18S rRNA* | ORIGENE | HP220445 |
| *GUCY2C* | ORIGENE | HP208194 |
| Recombinant DNA |  |  |
| Software and algorithms |  |  |
| Olympus |  | Olympus, Tokyo, Japan) |
| Fiji |  | ImageJ-2020 |
| Prism 6 and 8 | GraphPad | https://www.graphpad.com/scientific-software/prism/ |
| MACRO |  | ImageJ<sup>22</sup> |
| Other |  |  |
| Wnt3A Conditioned Medium (Atcc Crl-2647), | ATCC | Crl-2647 |
| Spondin1 Conditioned Medium | Gift From Calvin Kuo lab, Stanford University |  |
| Noggin Conditioned Medium | Gift From Gijs van den Brink Tytgat Institute for Liver and Intestinal Research |  |

## EXPERIMENTAL MODEL AND STUDY PARTICIPANT DETAILS

| Duodenal lines | Known Ailment | Gender | Ethnicity | Origin of biopsies |
| --- | --- | --- | --- | --- |
| Patient 1 | F508del mutation | NA | NA | Distal colon/Rectum |
| Patient 2 | F508del mutation | NA | NA | Distal colon/Rectum |
| Patient 3 | F508del mutation | NA | NA | Distal colon/Rectum |
| Patient 4 | F508del mutation | NA | NA | Ileum |
| Healthy subject 1 | normal | Female | NA | Distal colon/Rectum |
| Healthy subject 2 | normal | Male | Asian | Distal colon/Rectum |
| Healthy subject 3 | normal | Male | White | Distal colon/Rectum |
| Healthy subject 4 | normal | Male | Asian | Ileum |

823 NA=not available

## METHOD DETAILS

### Intestinal enteroid cultures from human biopsies

One ileal and three distal colon/rectal enteroids lines each from separate homozygous F508del-CF patients (adult) were obtained from Hugo R. de Jonge from the Biobank of Enteroids at the Hubrecht Foundation for Organoid Technology (MEC-2012-317). Normal human ileal and distal colonic/rectal enteroids lines established from 3 separate normal healthy subjects (NHS) including both male and female subjects were provided by the Hopkins GI Core Center. All experimental protocols were approved by the Johns Hopkins University Institutional Review Board (NA_00038329) and the Hubrecht Institute Human Research Review Board. Enteroid cultures were established from ileal and colonic biopsies obtained after endoscopic screening colonoscopies and upper endoscopies in normal subjects with no pathology found. Informed consent for tissue collection, generation, storage, and use of the enteroids was obtained from all participating patients. Enteroids were established and maintained as we have described ^15,57^. Briefly, enteroids generated from isolated intestinal crypts were maintained as cysts embedded in Matrigel (Corning, USA) in 24-well plates and cultured in Wnt3A, R-spondin, and Noggin-containing growth media, as we have described ^15^. The medium was refreshed every 2–3 d, and enteroids were passaged 1:3 every 7–10 d. Enteroids from passages 1–30 were used for these studies.

Homozygous F508del-CF and site-matched enteroids from separate NHS were studied in undifferentiated (UD), 3d-differentiated (PD), and 6d-differentiated (FD) states. Experiments were repeated at least 3-4 times and performed on each enteroid line.

### Forskolin-induced swelling of enteroids

Human enteroids from a 6-8-day-old culture were seeded in a flat-bottom 24-well culture plate in 10 μl Matrigel commonly containing 20–200 enteroids and 500 μl culture media. For studies on undifferentiated (UD) enteroids, the assays were performed 1 day after seeding. For partially (PD) and fully differentiated (FD) enteroids, the growth media were replaced with a differentiation media (without Wnt-3A and R-spondin) for 3 or 6 days respectively before the assay was performed. On the day of assay, growth or DF media were removed and enteroids were incubated for 60 min with 500 μl DMEM (without growth factors) containing 10 μM calcein green (Invitrogen). For optimal inhibition of transport proteins, enteroids were preincubated with CFTR_inh_-BPO (10 μM), DRAinh A250 (10µM), and CaCCinh-A01(15µM) (all gifts from Alan Verkman), dimethyl amiloride (2 µM) (Sigma-Aldrich, St. Louis, MO) for NHE1+2 inhibition, NHE3inh S3226 (20μM) (gift from Sinofi-Aventis). High-resolution spheroid images were captured using a Keyence BZ-X700 fluorescence microscope. Three wells were used to study one condition, and up to 60 wells were analyzed per experiment. For treatment with CFTR correctors, enteroids were preincubated for 24 h with tezacaftor (VX-661), and elexacaftor (VX-445) (3 µM each) (both from Selleck Chemicals LLC). For treatment with a CFTR potentiator, 5 μM VX-770 (Selleck Chemicals LLC) was added an hour before forskolin addition. The concentrations used were arbitrarily chosen since direct addition to the enteroids was assumed different from systemically administered drugs in patients.

Organoid swelling was measured as previously described ^11,22,44^. Enteroids were fluorescently labeled with calcein green and the total enteroid surface area per well over time was monitored by a Keyence BZ-X700 fluorescence microscope. Agonist-stimulated organoid swelling was calculated as described using an organoid swelling analysis macro for ImageJ ^22^. Briefly, to quantify swelling, total calcein-green surface areas were selected for each time point and expressed as a percentage increase with respect to the initial time point (T=0mins, set at 100%). The relative area increase was expressed per 15-minute time intervals and measurements were generated for each condition (T=0 to 60 min for Fsk or 120 min for linaclotide because the rate of secretion for linaclotide was slower than with cAMP). The area under the curve (AUC) (T=60 or 120 min, baseline 100%) was calculated using Prism.

### RNA isolation and Quantitative Real-Time PCR

Total RNA was extracted from 3D cultures using the PureLink RNA Mini Kit (Life Technologies, Carlsbad, CA) according to the manufacturer’s protocol. Complementary DNA was synthesized from 1 to 2 μg of RNA using SuperScript VILO Master Mix (Life Technologies). Quantitative real-time PCR was performed using Power SYBR Green Master Mix (Life Technologies) on a QuantStudio 12K Flex real-time PCR system (Applied Biosystems, Foster City, CA). Each sample was run in triplicate, and 5 ng RNA-equivalent complementary DNA was used for each reaction. Commercially available primer pairs from OriGene Technologies (Rockville, MD) were used: *LGR5* : HP207145; *Ki67* : HP206104; *MUC2* : HP206138; *CHGA* : HP205193; *DRA* : HP200096; *SI* : HP203589; *PDE4 :* HP233178; *18S rRNA* : HP220445. The relative fold changes in mRNA levels of genes were determined by using the 2 ^-ΔΔCT^ method with human 18S ribosomal, RNA simultaneously studied and used as the internal control for normalization and shown as fold change compared with the NHS-UD control.

### Adenoviral Flag-DRA, and cGKII infection

Triple Flag-tagged human DRA was cloned into the adenoviral shuttle vector ADLOX.HTM under the control of a cytomegalovirus promoter. The preparation of adeno-Flag-DRA and cGKII was reported previously ^33,58,59^. To infect with adenovirus (Adeno-Flag-DRA, or cGKII) enteroids were collected after melting the Matrigel and were mechanically dissociated through pipetting (30-50x). Enteroids were washed with basal media and centrifuged at 4^0^C, 900 x g for 5 min. The supernatant was discarded and the pellet was resuspended in enteroid growth media (500µl) combined with 10µl of the adenoviral solution with Y-27632 and CHIR99021(10µM) in one well of a 24-well plate. The plate was centrifuged at 32 °C, 600 x g, for 1 hr followed by incubation in a tissue culture incubator at 37^0^C at 5% CO_2_ for 6 hr. Adeno-infected enteroids were collected and centrifuged at 4^0^C, 900 x g for 5 min, and plated with Matrigel. The plate was incubated at 37 °C for 10 min to allow the Matrigel to polymerize, then a total of 500 μL of growth media with Y-27632 and CHIR99021 (10µM each) was added to the well. For the study, the Y-27632 and CHIR99021 containing growth media were replaced with UD or DF media as required for the assay.

### DRA knockdown in distal colonic/rectal enteroids

Enteroids were prepared as above and the adenoviral solution was replaced with a lentiviral solution containing 10mg/ml polybrene with 5µl control or DRA-shRNA + Y-27632 and CHIR99021(10µM each) in growth media (500µl). The shRNA-infected enteroids were collected and centrifuged at 4^0^C, 900 x g for 5 min, plated with Matrigel, and treated as above. Three days after the infection, enteroid media were replaced with growth media containing puromycin (0.2µg/ml) to maintain stable expression of the shRNAs.

### Immunofluorescence staining and confocal image analysis

Immunostaining of 3D enteroids was done in suspension. Briefly, recovered enteroids were fixed in 4% paraformaldehyde in 10 mmol/L phosphate buffer (pH 7.4) for 30 minutes at 4°C and then washed 2× with phosphate-buffered saline. Enteroids were permeabilized and stained in phosphate-buffered saline with 2% bovine serum albumin, 15% fetal bovine serum 0.25% Triton X-100, and 1% saponin for 60 minutes (all Sigma-Aldrich St. Louis, MO, USA), followed by overnight incubation with antibodies. Primary antibodies used: mouse anti-MUC2 (sc7314, Santa Cruz Biotechnology, Dallas, TX), anti-E-cadherin (Clone 36, BD Biosciences) anti-GUCY2C (LS-C498735, LSBio, Lynnwood, WA), anti-ChgA (20085, Immunostar). All primary antibodies were diluted at 1:100 and incubated overnight. Stained cells were then washed 3 times for 10 min each with PBS followed by incubation with Alexa-488/568 conjugated Goat anti-Rabbit and anti-mouse antibodies respectively (Molecular Probes/Invitrogen, USA) diluted 1:500 in PBS. Hoechst (Vector Laboratories, USA) was used at a 1:1000 dilution in PBS for nuclear/DNA labeling. After incubation, cells were washed 3 times for 10 min each and mounted in ProLong Gold (Vector Laboratories, USA) overnight at 4 °C. Images were collected using ×20 objective on FV3000 confocal microscope (Olympus, Tokyo, Japan) with Olympus and ImageJ software (NIH).

### Immunoblotting

For protein detection, 3D enteroids from NHS or F508del-CF patients were harvested in phosphate-buffered saline after melting the Matrigel and rinsed 2 times with phosphate-buffered saline. Cell lysate preparation and Western blot were performed using validated antibodies for DRA (SC-376187; mouse monoclonal, 1:500; Santa Cruz, Dallas, TX), <u>CFTR (CF660, CF217 from CF consortium), and anti-CFTR mAb 596 (nucleotide-binding domain 2 [NBD2] 1204–1211 aa) (Cystic Fibrosis Foundation,</u> https://www.cff.org<u>), diluted 1:500 in TBST 2% (w/v) skim milk cGKII (SC-393126; mouse monoclonal, 1:500; Santa Cruz,</u> Dallas, TX) and glyceraldehyde-3-phosphate dehydrogenase (G8795; mouse monoclonal, 1:5000; Sigma-Aldrich). Protein bands were visualized and quantitated using an Odyssey system and Image Studio software (LI-COR Biosciences, Lincoln, NE).

### Single-cell RNA sequencing analysis of normal human small intestine

The entire small intestine from two deceased but otherwise, normal people were obtained from the research-consent deceased organ donor program obtained through an IRB-approved research protocol with donor Network West in collaboration with the UCSF Viable Tissue Acquisition Lab Core. The studies with this tissue were described previously that include single-cell RNA-sequencing of the entire small intestine divided along the horizontal axis into 30 segments, which for the current report were incorporated into a single analysis consisting of the entire small intestine ^52^. As reported, villus zonation scoring used MATLAB version 2018b to annotate the enterocytes according to their position along the crypt-villus axis, ^60^. In addition, villus domains were assigned using landmark genes to separate villus enterocytes into 6 zones.

## QUANTIFICATION AND STATISTICAL ANALYSIS

Statistical testing based on the raw data was assessed using Prism statistical software version 6 and 8 (GraphPad). Significance was considered *p* < 0.05 using Student’s t-tests or One-way ANOVA with Bonferroni post hoc test. All experiments were performed on 3 different colonoid lines from healthy subjects and CF-F508del-CF patients. Some experiments were also repeated on 1 ileal enteroid line from a healthy subject and a F508del-CF patient. N refers to the number of independent replicates performed.

## Supplementary Figures

**Supplementary Figure 1.**
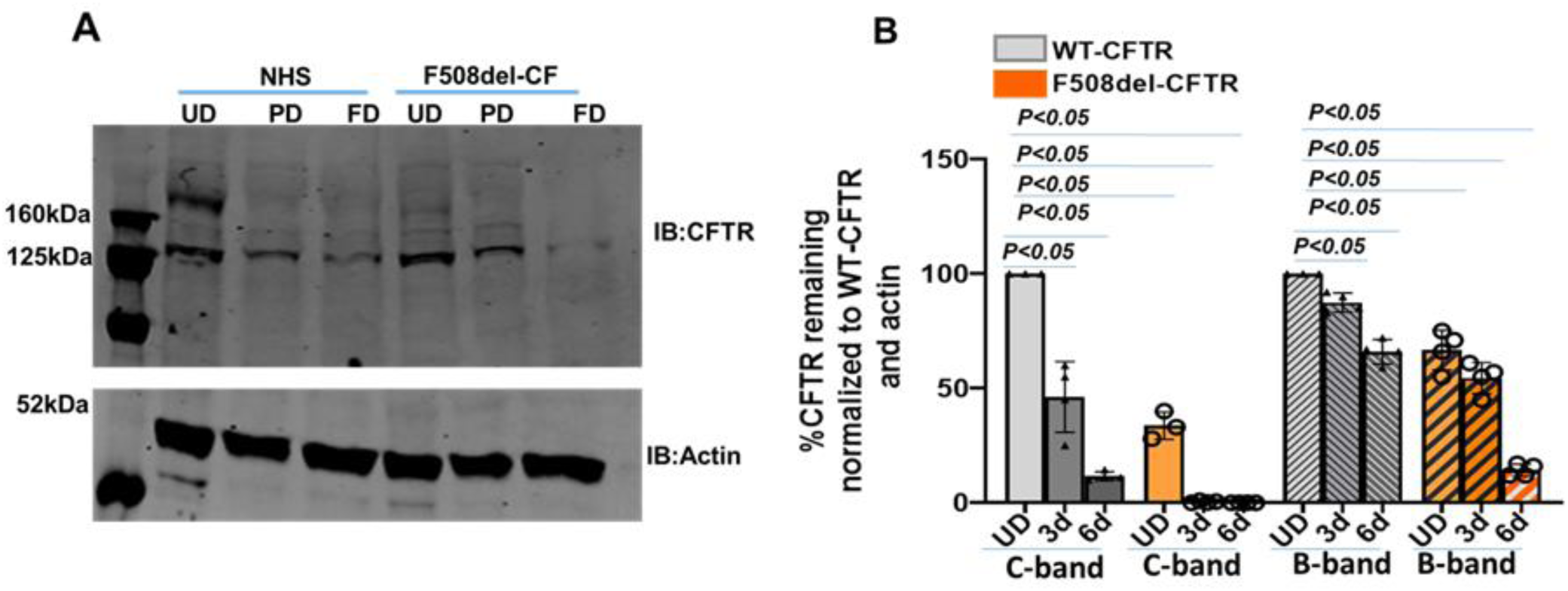
CFTR expression decreases with the differentiation colonic/rectal enteroids. **(A)** A representative western blot showing CFTR expression in human colonic/rectal enteroids in undifferentiated (UD), partially differentiated (PD) and differentiated (FD) states. **(B)** Quantitation of multiple blots showing a decrease in CFTR C and B band expression with differentiation of enteroids. The bar graph represents % change in C and B band expression compared to WT-CFTR normalized to loading control β-actin (means ± SEM). Analysis of differences was determined with a one-way ANOVA. Data points represent an average of 3-4 independent assays from 3 NHS and 3 CF patients.

**Supplementary Figure 2.**
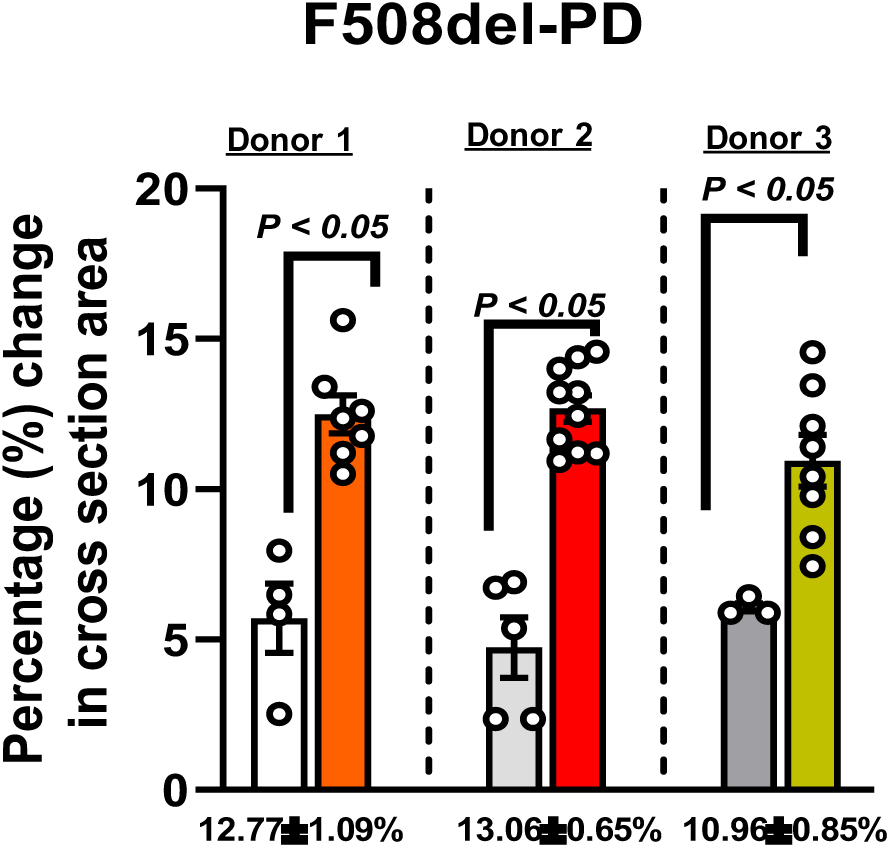
FIS response in PD colonic/rectal enteroids in 3 separate F508del CF patients. Results are calculated as the percentage change in surface area relative to t = 0 (percentage change in cross-section area) measured at 10-15-min time intervals for 1 h. The bar graph depicts fluid secretion plots (t = 60 min, means ± SEM). Data points represent an average of 3-5 independent assays each containing 50-100 enteroids from 1 CF patient. P value was determined using Student’s t-test.

**Supplementary Figure 3.**
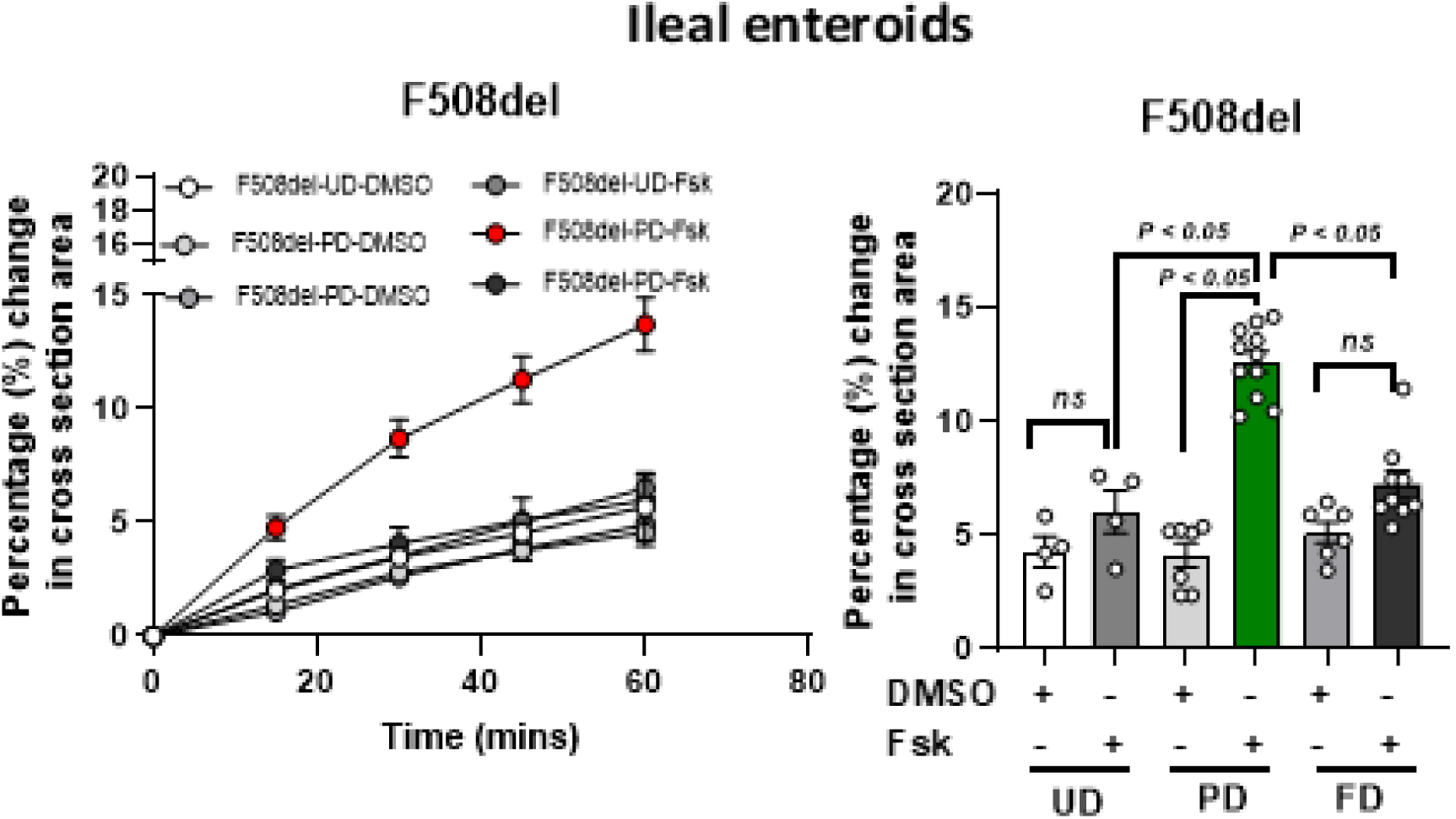
FIS response in UD, PD, and FD ileal enteroids from a F508del CF patient. Results as the percentage change in surface area relative to t = 0 measured at 15-min time intervals expressed for 1 h (means ± SD), and the bar graph depicts fluid secretion plots (t = 60 min, means ± SEM). An average of 3-6 swelling assays each containing 50-100 enteroids from a F508del CF patient is shown. Analysis of differences was determined with a one-way ANOVA and Bonferroni post hoc test, ns=not significant.

**Supplementary Figure 4.**
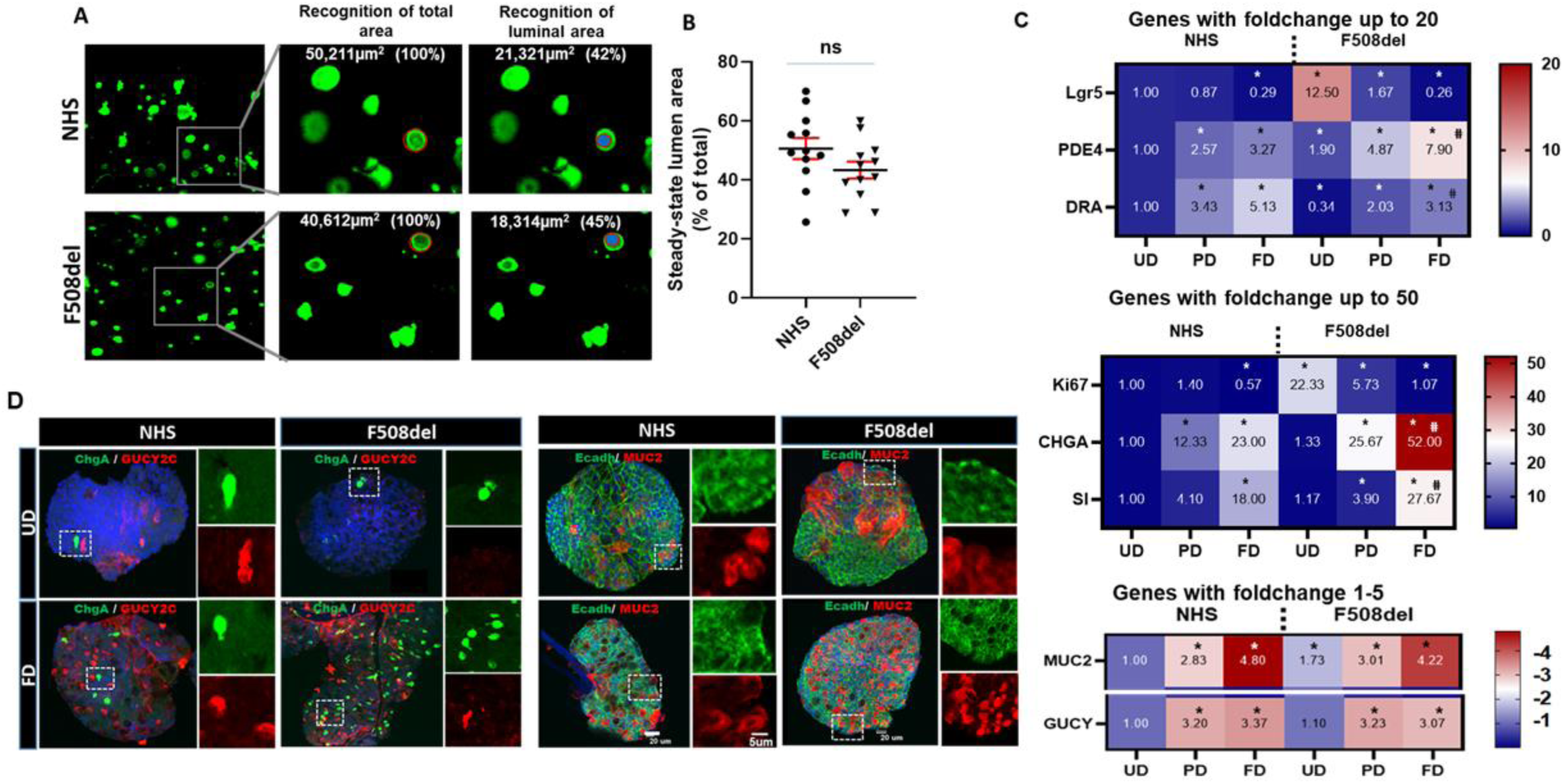
F508del-CF enteroids undergo differentiation similarly to the enteroids from normal healthy subjects (NHS) and have similar Steady State Lumen Area (SLA). **(A)** Recognition of the total lumen area of colonic/rectal enteroids from NHS and F508del (*xy* plane) determined by Volocity imaging software. The luminal surface area (μm^2^ and %) of the total enteroids surface area (100%) is indicated at the top of the image. **(B)** Quantification of the SLA of enteroids (luminal area/total area) from NHS (*N =3*) or F508del (*N=3*) is shown. Four independent experiments were performed for each subject. ns=not significant by Student’s unpaired t-test. **(C)** Changes in mRNA expression of selected genes were determined by qRT-PCR and relative fold change was calculated between UD (set as 1), PD, and FD enteroids normalized to *18S* ribosomal RNA as the endogenous control. The mean ± SEM of fold change was calculated and is represented as a heatmap. N= 3-4 independent analysis on enteroids from a single individual. * denote *P<0.05* compared to UD NHS enteroids. **^#^** ^d^enotes *P<0.05* compared to FD NHS enteroids. **(D)** Representative immunofluorescence staining showing expression of CHGA (green), and GUCY2C (red) (left); E-cadh (green) and MUC2 (red) (right) in UD (above) and FD (below) NHS and F508del colonic/rectal enteroids. An XY optical section of a multi-z-stack confocal section (1µm) is shown. Scale bar: 20µm. The insets display a magnification of the area in the white box. Scale bar: 5µm.

**Supplementary Figure 5.**
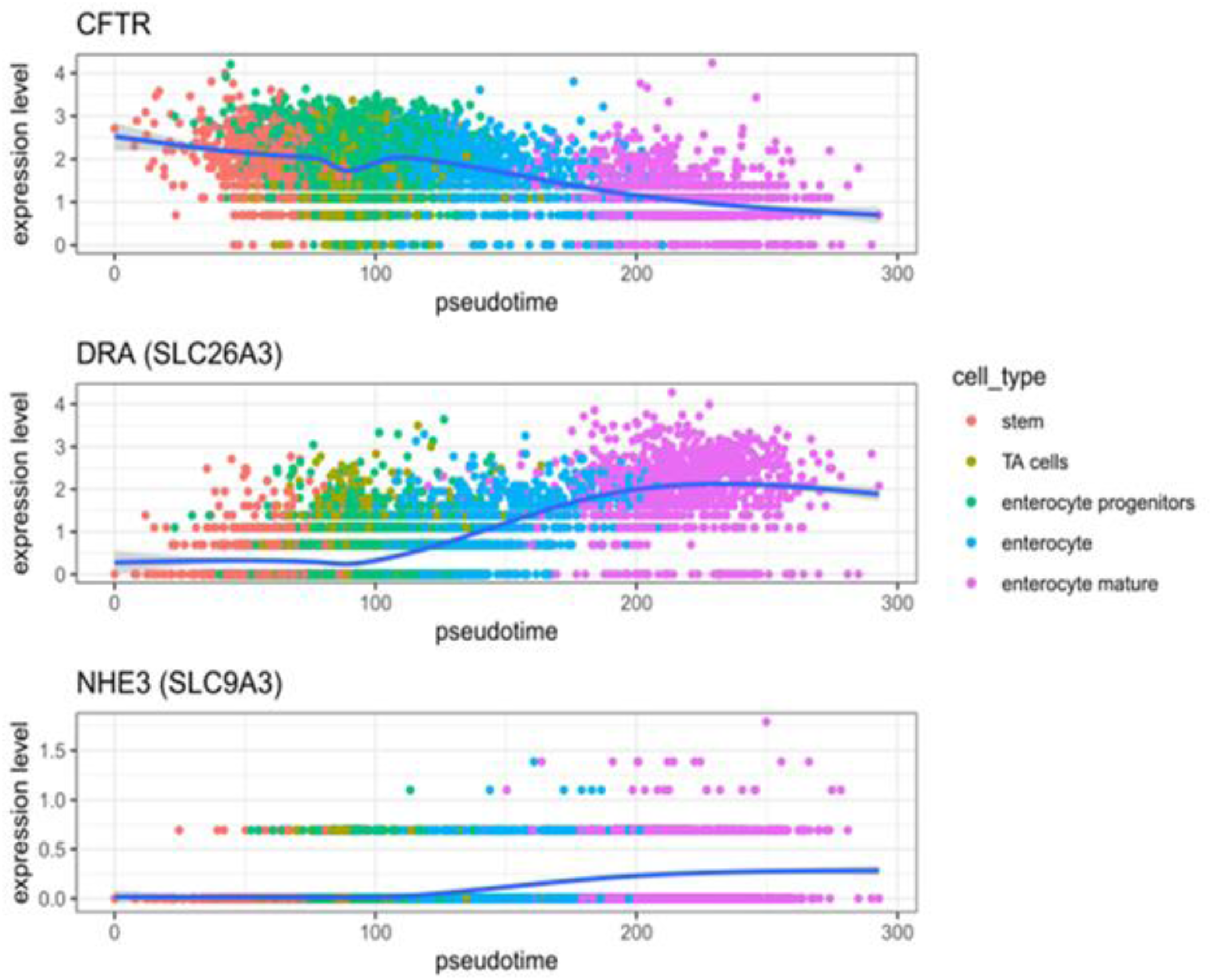
Single-cell RNAseq analysis for important ion transporters from the entire small intestine of one donor. **S**ingle-cell RNA sequencing was reported previously on normal human small intestine^55^. Reanalysis of sc-RNAseq for important ion transport proteins was performed by determining the differentiation pseudotime of cells of the absorptive lineage. Shown here is the expression of the CFTR, DRAand NHE3 genes as a function of differentiation pseudotime. Results suggest changes in mRNA expression of CFTR, DRA, and NHE3 that occurred in the transition from enterocytes to mature enterocytes (150-200) were similar to the protein expression changes that occur in PD enteroids. The cells in this transition zone are considered representative of the PD enterocyte population.

**Supplementary Figure 6.**
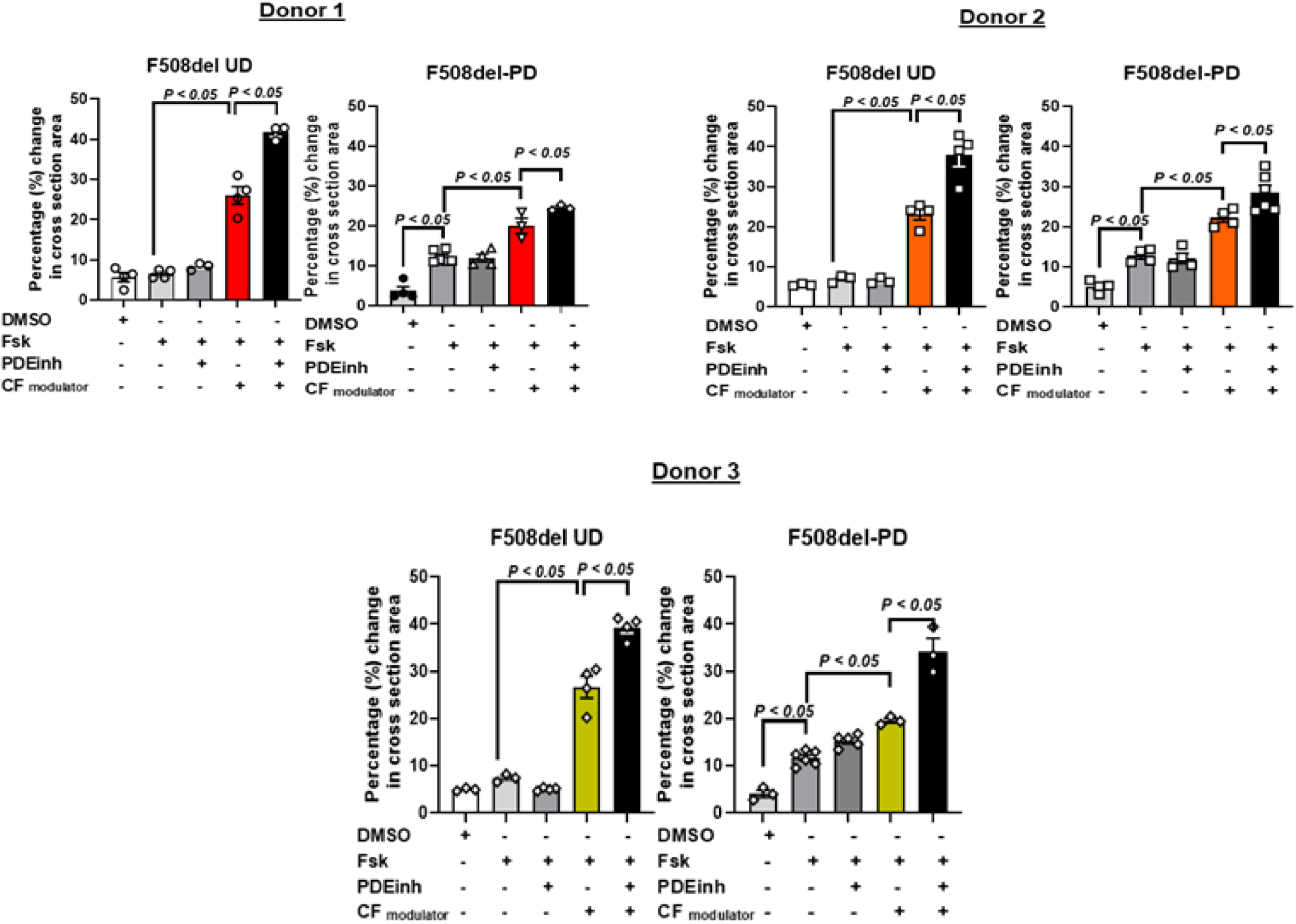
Theophylline enhances FIS response from CFTR-modulators-treated PD enteroid in 3 separate F508del CF patients. Effect of theophylline (100µM-1h) or combined CFTR-modulators (tezacaftor:VX-661+elexacaftor:VX-445 3µM each for 24h; ivacaftor:VX-770 (5µM) for 1h) and in combination on FIS (5µM) response in UD, and PD F508del distal colonic/rectal enteroids from donor 1, donor 2 and donor 3. Results are calculated as the percentage change in surface area relative to t = 0 measured at 15-min time intervals for 1 h and fluid secretion is represented as bar graph plots (t = 60 min, means ± SEM). An average of 3-6 swelling assays each containing 100-200 enteroids from each F508del CF patient is shown separately. Analysis of differences was determined with a one-way ANOVA and Bonferroni post hoc test.

**Supplementary Figure 7.**
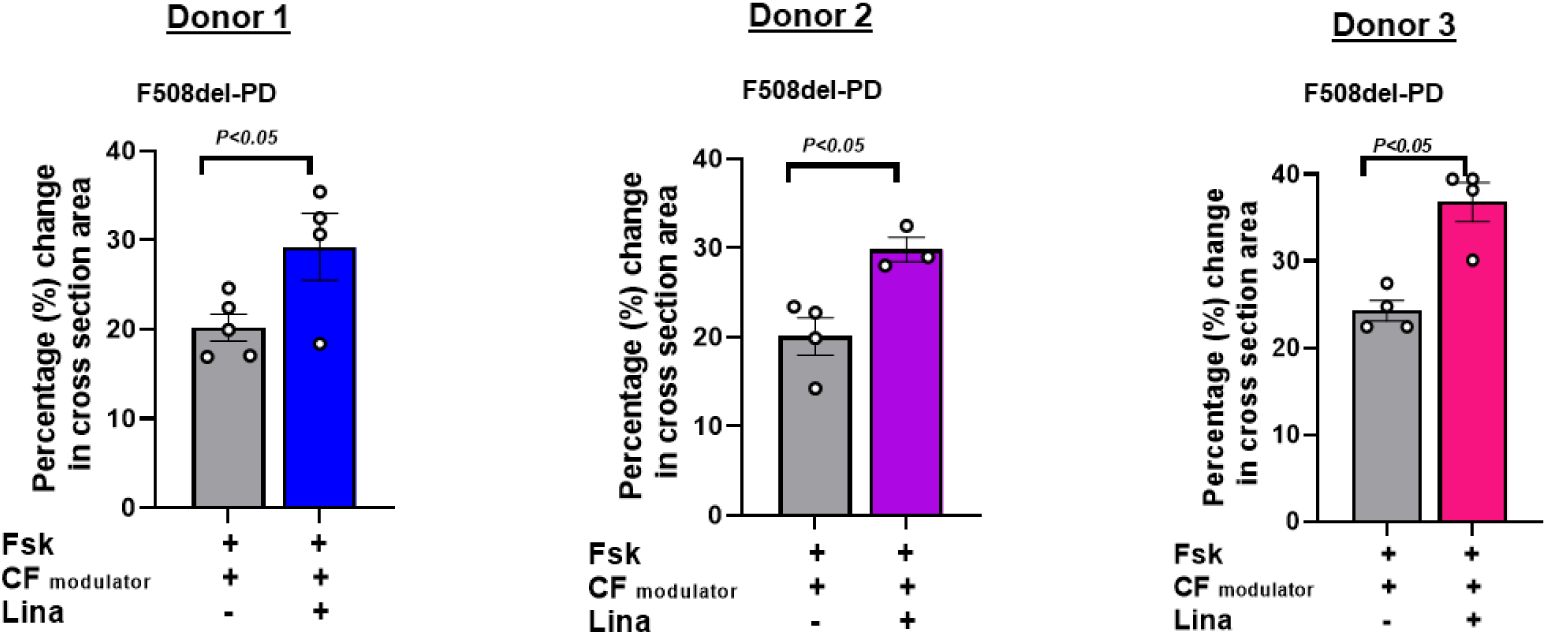
Linaclotide enhances combined CFTR-modulator-mediated FIS response from PD colonic/rectal enteroids in 3 separate F508del CF patients. PD F508del enteroids pretreated with tezacaftor:VX-661+elexacaftor:VX-445 3µM each for 24h, and ivacaftor:VX-770 (5µM) for 1h were used to determine FIS in response to Fsk (5µM) or with Linaclotide (10µM). A time-dependent increase in organoid surface area normalized to that at t = 0 was measured at 15-minute time intervals for 2 h and represented as a bar graph depicting fluid secretion plots (t = 120 min, means ± SEM). Data points represent an average of 3-5 independent assays each of 50-100 enteroids done separately in each CF patient. Data are means ± SEM with *P* value using Student’s t-test.

**Table 1.** Single-cell RNAseq analysis of human intestinal villus distribution of intestinal transport proteins. Reanalysis of sc-RNAseq results from human small intestine suggests changes in mRNA expression of DRA, NHE3, and CFTR that occurred in the differentiation pseudotime interval 150-200, which roughly corresponds to the transition between enterocytes and mature enterocyte villus zones from the villus base (see Figure S4) that was similar to the PD enteroids. We suggest that cells in this transition zone are representative of the PD enterocyte population. The upper part of the Table is the number of cells in each pseudotime interval that express CFTR, SLC26A3, and SLC9A3 and their combinations. The bottom part of the Table is the percent of cells in each pseudo time interval expressing the specific transporters.

| pseudotime interval | number of cells | CFTR | SLC26A3 | SLC9A3 | CFTR_DRA | CFTR_NHE3 | DRA_NHE3 | CFTR_DRA_NHE3 |
| --- | --- | --- | --- | --- | --- | --- | --- | --- |
| (100,150] | 1872 | 1840 | 991 | 88 | 975 | 88 | 64 | 64 |
| (150,200] | 498 | 463 | 472 | 142 | 438 | 133 | 132 | 123 |
| (200,293] | 782 | 706 | 745 | 263 | 669 | 225 | 247 | 209 |

**Table 1. Single-cell RNAseq analysis of human intestinal villus distribution of intestinal transport proteins.**
| pt_bin | all.cells | CFTR | SLC26A3 | SLC9A3 | CFTR_DRA | CFTR_NHE3 | DRA_NHE3 | CFTR_DRA_NHE3 |
| --- | --- | --- | --- | --- | --- | --- | --- | --- |
| (100,150] | 100.0% | 98.3% | 52.9% | 4.7% | 52.1% | 4.7% | 3.4% | 3.4% |
| (150,200] | 100.0% | 93.0% | 94.8% | 28.5% | 88.0% | 26.7% | 26.5% | 24.7% |
| (200,293] | 100.0% | 90.3% | 95.3% | 33.6% | 85.5% | 28.8% | 31.6% | 26.7% |

